# Fate of developmental mechanisms of myocardial plasticity in the postnatal heart

**DOI:** 10.1101/842864

**Authors:** Konstantinos E. Hatzistergos, Michael A. Durante, Krystalenia Valasaki, J. William Harbour, Joshua M. Hare

## Abstract

Whether the heart’s organ-founding, progenitor cell gene regulatory networks (CPC-GRNs) are sustained after birth and can be therapeutically evoked for regeneration in response to disease, remains elusive. Here, we report a spatiotemporally resolved analysis of CPC-GRN deployment dynamics, through the pan-CPC-GRN gene *Isl1*. We show that the Isl1-CPC-GRNs that are deployed during early cardiogenesis and generate the cardiomyocyte majority from mesoderm, undergo programmed silencing through proteasome- and PRC2-mediated Isl1 repression, selectively in the arterial pole. In contrast, we identify a neural crest (CNC)-specific Wnt/β-catenin/Isl1-CPC-GRN that is deployed through the venous pole during cardiac growth and partitioning, and contributes a minority of cardiomyocytes which, in turn, expand massively to build ~10% of the biventricular myocardium. These “dorsal CNCs” continue to sporadically generate cardiomyocytes throughout postnatal growth which, however, are non-proliferative, suggesting that partitioning-like, fetal proliferation signals could be therapeutically targeted to evoke clonal expansion capacity in postnatal CNC-cardiomyocytes for heart regeneration.

## INTRODUCTION

The four-chambered, mature, adult mammalian heart forms through a series of complex, spatially and temporally controlled, morphogenetic processes which evolve during embryogenesis. One of the earliest organ-founding processes is the activation of a cardiac progenitor cell (CPC) gene regulatory network (GRN) expressing the transcription factor *Isl1*, in the anterior lateral plate mesoderm (Cai et al., 2003; Prall et al., 2007). The splanchnic layer of this region will produce the two main heart lineages, the first (FHF) and second (SHF) heart fields, from which the majority of the beating cardiomyocytes (CMs) will differentiate (Cai et al., 2003; Prall et al., 2007).

The FHF is specified through mechanisms that lead to the inactivation of the Isl1-CPC-GRN and activation of a CPC-GRN expressing the master cardiac transcription factor Nkx2-5 (Lyons et al., 1995; Prall et al., 2007). In turn, the FHF primordia differentiate into a linear, beating heart tube which is patterned to an arterial and venous identity along its anteroposterior axis, and will later morph into the left ventricle (LV), parts of the atria and the atrioventricular canal (Lyons et al., 1995).

The SHF forms medially and dorsally to the FHF, from the splanchnic mesoderm region ventrally to the pharynx, and becomes specified into an anterior (aSHF) and posterior (pSHF) domain, through mechanisms that involve sustained Isl1-CPC-GRN activity (Cai et al., 2003; Prall et al., 2007). Importantly, the aSHF and pSHF Isl1-CPC-GRNs are activated through different *Isl1 cis*-regulatory elements (Kang et al., 2009; Kappen and Salbaum, 2009). The aSHF-Isl1-GRN is activated together with the Nkx2-5-CPC-GRN and deploys CPCs through the arterial pole of the heart tube, which will generate the right ventricle (RV), parts of the interventricular septum (IVS) and outflow tract (OFT) (Cai et al., 2003). The pSHF-Isl1-GRN deploys a mixture of Nkx2-5 expressing and non-expressing CPCs through the venous pole which will give rise to the dorsal mesocardium, pulmonary and caval veins as well as parts of the cardiac conduction system and atria (Meilhac and Buckingham, 2018). Continuous recruitment of CPCs from the arterial and venous poles through the aSHF-and pSHF-Isl1-GRNs respectively, causes the linear heart tube to grow, elongate and bend, and eventually undergo a rightward looping which shifts its venous and arterial poles to their definitive positions at the ventral (outflow) and dorsal (inflow) domains of the cardiac base, respectively (Meilhac and Buckingham, 2018).

The arterial and venous poles are now oriented along the dorsoventral rather than anteroposterior embryonic axis, and the heart enters its growth and partitioning phase. A layer of trabecular myocardium grows along the endocardial surface of the ventricles which later undergoes compaction and expands clonally into IVS myocardium (Meilhac and Buckingham, 2018; Meilhac et al., 2003). Interestingly, Isl1-GRN perturbations produce myocardial trabeculation, septation and compaction defects, suggesting important roles of Isl1-GRNs in these processes (Cai et al., 2003; Cho et al., 2018; Ma et al., 2008). Concurrently, neural ectoderm-derived cardiac neural crest cells (CNCs) are deployed from the rhombencephalon (Kirby et al., 1983). Particularly, an early wave of CNCs migrates through the pharyngeal arches into the arterial pole to primarily contribute to the aorticopulmonary septation and smooth muscle cells (SMCs), as well as parasympathetic innervation (Meilhac and Buckingham, 2018); whereas a second wave of CNCs are thought to migrate later through the pSHF-derived dorsal mesocardium into the venous pole region, to contribute sympathetic nerves and endocardial cushion mesenchyme (Hildreth et al., 2008; Nakamura et al., 2006; Verberne et al., 1998). Importantly, recruitment of CNCs through the arterial and venous pole regions are also mediated by Isl1-GRNs (Engleka et al., 2012; Sun et al., 2008; Sun et al., 2007).

In mammals, these organ-founding CPC-GRNs are thought to be silenced before birth, leaving the postnatal heart unable to regenerate muscle damages (Li et al., 2019; Meilhac and Buckingham, 2018). However, some studies indicate that, in rodents and humans, the aSHF Isl1-CPC-GRN is sustained in the arterial pole region after birth and could be therapeutically targeted for postnatal heart regeneration (Laugwitz et al., 2005; Meilhac and Buckingham, 2018). Accordingly, here, we investigate this possibility through a combination of *Isl1*-targeted, multicolor lineage-tracing, functional mutagenesis and single-cell RNA sequencing (scRNAseq) experiments. Our results show that the early cardioblastic Isl1-CPC-GRNs, which produce the FHF and SHF as well as the CNCs which are recruited through the arterial pole, undergo programmed silencing through selective activation of the ubiquitin/proteasome and PRC2 pathways in the outflow region of the heart tube and, therefore, are unlikely to be sustained in the postnatal heart. However, we provide evidence that the Isl1-GRN which recruits CNCs through the venous pole during partitioning of the heart tube, is sustained in the inflow region throughout the fetal and postnatal growth phases via a Wnt/β-catenin/Isl1 feedback loop. We further show that, in addition to sympathetic nerves and cushion mesenchyme, these “dorsal CNCs” produce a small number of biventricular trabecular CMs which undergo massive clonal expansion during partitioning and growth of the fetal heart, to eventually produce ∼10% of the postnatal biventricular myocardium. After birth, this dorsal Isl1-CNC-GRN continues to sporadically generate CMs which however are restricted from undergoing clonal expansion; suggesting that activation of partitioning-like fetal proliferation cues might provide a novel therapeutic approach to enhance the production of regenerative CNC-CMs in the adult heart in response to damage.

## RESULTS

### Diverse developmental origins and identities of postnatal cardiac cells with sustained *Isl1* activity

To monitor the expression of the pan-CPC-GRN gene *Isl1*, we employed a previously described mouse line carrying a *loxP*-flanked *Isl1-nLacZ* allele [Fig. S1A] (Sun et al., 2007). This *Isl1-nLacZ* knock-in reports *Isl1* transcription from both neural (Bejerano et al., 2006; Uemura et al., 2005) and mesoderm-specific (Kang et al., 2009; Kappen and Salbaum, 2009) regulatory elements. Accordingly, as previously shown (Sun et al., 2007), *Isl1-nLacZ* is expressed in aSHF and pSHF CPCs during early cardiogenesis [Figure S1B] and later in CNC cells [Figure S1C]. After birth, its cardiac expression is sustained in 5 distinct dorsoventral (DV) domains [Fig. S1D-E]. Ventrally (outflow domain), *Isl1-nLacZ* is expressed in cells within the proximal aorta (Ao) and pulmonary artery (PA), as well as the OFT myocardium [Fig. S1D]. Dorsally (inflow domain), *Isl1-nLacZ* labels cardiac ganglia (CGs) and the sinoatrial node (SAN) [Fig. S1E].

The ventral domain of the cardiac base develops from the arterial pole of the heart tube through the FHF, aSHF-and CNC Isl1-CPC-GRNs (Engleka et al., 2012; Sun et al., 2007); whereas, the dorsal domain develops from the venous pole through the FHF, pSHF (including the SAN) and CNC Isl1-CPC-GRNs (Hildreth et al., 2008; Nakamura et al., 2006; Sun et al., 2007; Verberne et al., 1998). Therefore, to determine the Isl1-CPC-GRNs through which the ventral and dorsal *Isl1-nLacZ* activities are sustained after birth, we interbred the *Isl1-nLacZ* to the aSHF-specific *MEF2c-AHF-Cre (Verzi et al., 2005)* and CNC-specific *Wnt1-Cre2 (Lewis et al., 2013)* mouse lines. Under this strategy, the *loxP*-flanked *Isl1-nLacZ* reporter is conditionally excised from CNC- and aSHF-derived heart cells, respectively. Consequently, expression of *Isl1-nLacZ* persists in non-recombined cells [Fig. S2A].

As expected, in *Isl1-nLacZ;Mef2c-AHF-Cre* neonatal hearts (*nLacZ* excision from aSHF) the dorsal x-gal signal is not affected, indicating that the dorsal *Isl1-nLacZ^+^* domains are not sustained through the aSHF-Isl1-CPC-GRN. However, there is a ∼7-fold reduction (*p*<0.0005) in the ventral x-gal signal, indicating that most, but not all, of the ventral *Isl1-nLacZ* activity is sustained through the aSHF-Isl1-CPC-GRN. Immunohistochemical analysis determined that the remaining x-gal^+^ ventral cells are mostly SM22a^+^ SMCs and tyrosine hydroxylase (TH)^+^ sympathetic neurons within the proximal PA and Ao. Surprisingly, few α-sarcomeric actinin^+^/ x-gal^+^ CMs within the OFT base are also present, indicating that non-aSHF-Isl1-CPC-GRNs contribute a minority of ventral *Isl1-nLacZ^+^* CMs in this region [Fig. S2B-C and S3 A-E].

Comparably, in *Isl1-nLacZ;Wnt1-Cre2* neonatal hearts (*nLacZ* excision from CNCs), the ventral x-gal signal is minimally reduced, indicating that only a small proportion of the ventral *Isl1-nLacZ* activity is sustained through the Isl1-CNC-GRN [Fig. S2F-G, N]. Dorsally, it remains unchanged in the SAN but, as expected, extinguished from the CNC-derived CGs [Fig. S2H-I, N]. Interestingly, in triple-mutant *Isl1-nLacZ;Wnt1-Cre2;Mef2c-AHF-Cre* mice (*nLacZ* excision from both aSHF and CNCs), x-gal is sustained in few sm22a^+^ SMCs in the proximal PA and the dorsolateral wall of the Ao, but is extinguished from TH^+^ neurons and α-sarcomeric actinin^+^ CMs [Figs. S2J-K, N, and S3F]; whereas dorsally, x-gal remains remains unaffected in the SAN but extinguished from CGs [Figs. S2L-N, S3G].

Collectively, these *Isl1-nLacZ* lineage-tracing experiments delineate that sustained activity of the pan-CPC-GRN gene *Isl1* in the venous pole-derived, dorsal domain of the postnatal heart, identifies CNC-derived cells and the SAN. Whereas, sustained *Isl1* activity in the arterial pole-derived, ventral domain of the postnatal heart identifies a mixture CNC-derived TH^+^ neurons; OFT base CMs (most of which derive from the aSHF and some from the CNC); as well as SMCs in the proximal PA and Ao, which descend from at least 3 distinct developmental origins-the aSHF, CNC and a third, venous pole-derived lineage.

### Ventral but not dorsal *Isl1* is transcriptionally silenced

By PN week 4, *Isl1-nLacZ* activity in the arterial pole-derived, ventral domain of the postnatal heart has diminished [Fig. S1D, F, H]. In particular, x-gal signal is reduced in the PA and virtually extinguished from the OFT base and Ao, except from few cells in the dorsolateral Ao wall medially to the SAN [Fig. S1D, F, H, J]. No remarkable changes are observed in the dorsal (SAN and CG) *Isl1-nLacZ* activity [Fig. S1E, F-I].

To understand why the ventral but not dorsal Isl1-CPC-GRNs are progressively lost after birth, the dorsal (including the SAN and CGs) and ventral (including the Ao, PA and OFT base) domains of the cardiac base were dissected from *Isl1^+/+^* neonatal mice, dissociated into single-cell suspensions, recovered in culture for 5 days, and processed for single-cell RNA sequencing (scRNAseq) [Figure 1A]. A total of 11,418 ventral and 9,893 dorsal cells were sequenced. Analysis of transcriptional identities by Uniform Manifold Approximation and Projection (UMAP) dimensional reduction using Seurat (Stuart et al., 2019) generated a total of 15 transcriptionally and spatially distinct clusters [Figure 1B-D, and Table S1]. To classify the clusters, we first performed an unbiased gene set enrichment analysis of cluster-specific genes [Table S1] (Chen et al., 2013) and the resulting cell types were elaborated further by profiling the expression of 48 canonical marker genes [Figure 1C-D]. Accordingly, cluster 0 is enriched in *Sox9*, *Fbn1* and *Nfatc1* and is therefore assigned a cardiac valve cell identity. Clusters 1, 2 and 5 are enriched in *Fap* and are therefore classified as fibroblasts. Cluster 3 is enriched in Wt1 and is therefore assigned an epicardial identity. Clusters 4, 6 and 8 are enriched in *Myh11 and Tagln*, and are therefore classified as smooth muscle cells. Cluster 7 is classified as CNC-derived cells based on the expression of *Sox2, Sox10, Plp1, Nes, Ngfr, Foxd3* and *Tfap2a.* Cluster 9 is enriched in *Pecam1, Cdh5, Kit*, *Kdr*, and *Nr2f2* and is therefore classified as venous endothelium. Cluster 10 is assigned a working myocardial cell identity based on the expression of *Nkx2-5, Myh6, Myh7, Ttn, Tnni3, Tnnt2, Myl7* and *Nppa*. Cluster 11 is identified as low quality/dying ventricular CMs due to high mitochondrial contamination [Fig. S4A] (Stuart et al., 2019). Clusters 12 and 13 are classified as immune cells (B/T-lymphocytes and macrophages, respectively); and finally, cluster 14 is identified as conduction system CMs (which include the dorsal cluster of *Nkx2-5* negative/ *Isl1^+^* sinoatrial node CMs), based on the expression of *Isl1, Gja5, Ttn, Tnnt2* and *Myh7.* Interestingly, none of the clusters showed enrichment for *Isl1* expression [Figure 1C-E]. Rather, *Isl1*-expressing cells were dispersed in 14/15 clusters revealing a remarkable diversity of postnatal *Isl1^+^* cardiac cells [Figs. 1E and S4B].

**Figure 1.**
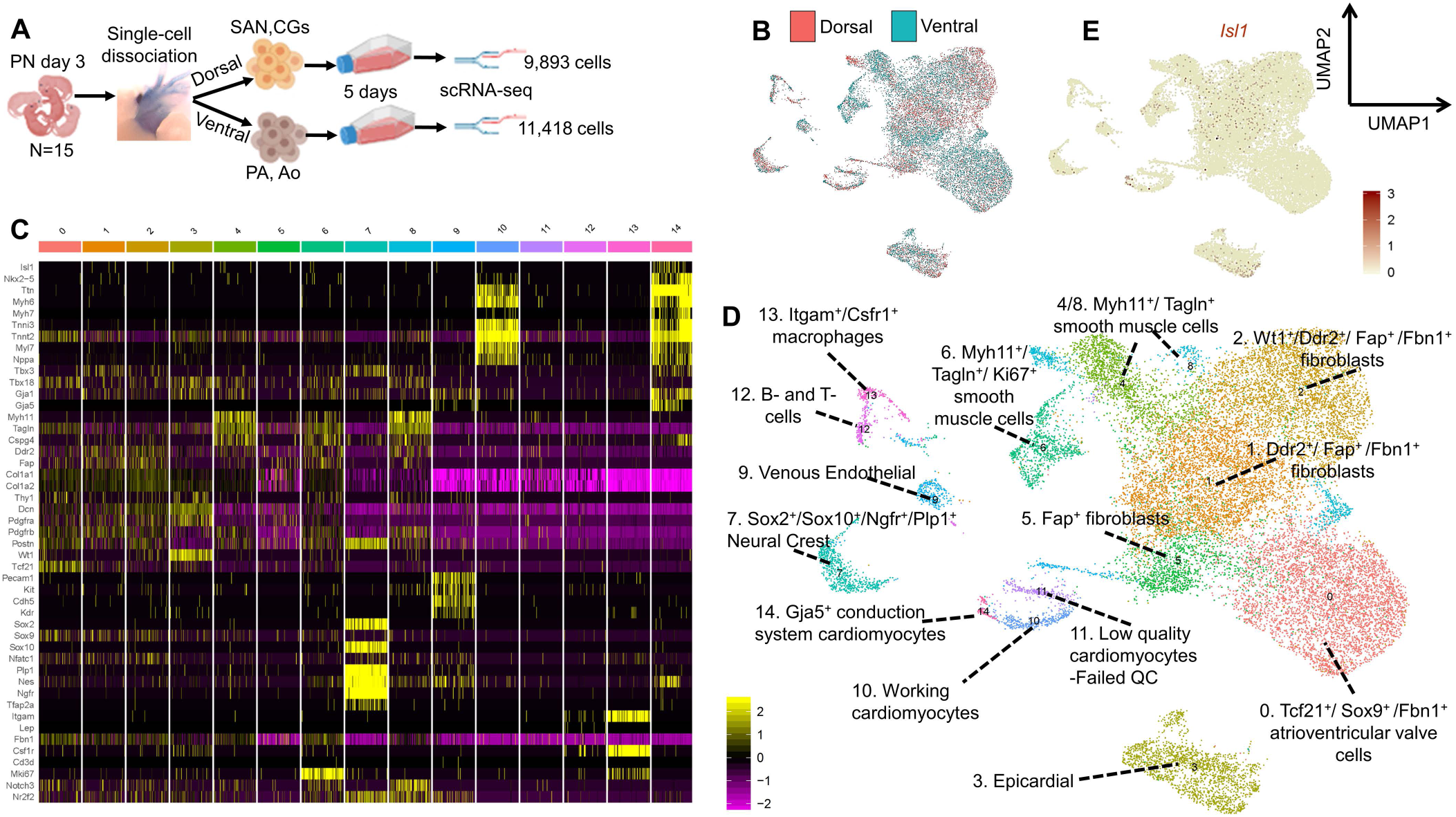
scRNAseq of arterial (ventral) and venous (dorsal) pole-derived neonatal cells. A, Experimental outline. **B**, Uniform Manifold Approximation and Projection dimensional reduction (UMAP) plot of 21,311 scRNAseq profiles following integration of the postnatal (PN) day 3 dorsal and ventral datasets. **C**, Heatmap of 47 lineage-specific genes across the 15 UMAP clusters. For visualization, each cluster is downsampled to 100 cells. **D**, UMAP plot of classified cell clusters. **E**, Expression of Isl1, visualized on UMAP plots. PA, pulmonary artery; Ao, aorta; CGs, cardiac ganglia; SAN, sinoatrial node.

Next, the *Isl1^+^* cells were subselected from the scRNAseq datasets based on their DV identities and subjected to differential gene expression (DEG) analysis [Fig. 2A]. Overall, there were 278 ventral and 205 dorsal *Isl1*^+^ cells with 567 DEGs [Fig. 2B, Table S2]. Unsupervised hierarchical clustering ordered the z-score normalized DEGs according to the DV identities of *Isl1^+^* cells [Fig. S5]. Functional enrichment analyses mapped multiple processes to the *Isl1*^+^ DEGs [Fig. 2C; Tables S2-S3], including processes related to heart and CNC development, regulation of transcription, as well as PRC2 (polycomb repressive complex 2)-related (Suz12 and Ezh2) gene silencing [Fig. 2D; Tables S3-S4]. Based on these findings we hypothesized that loss of ventral *Isl1-nLacZ* activity is driven by PRC2-mediated transcriptional silencing of the *Isl1* gene. To functionally test this possibility, cells from the ventral *Isl1* domain of PN week 4 *Isl1-nLacZ* hearts (which have already lost *Isl1-nLacZ* activity, i.e. Fig. S1) were cultured for 5 days in the presence or absence of the PRC2-specific, small molecule inhibitor EED226 [Fig. 2E] (Qi et al., 2017). Western blot and immunocytochemical analyses confirmed the successful inhibition of PRC2 activity in EED226-treated cells, as indicated by downregulation of H3K27me3 expression compared to controls [Fig. 2F]. Gene-expression and x-gal analyses demonstrated that inhibition of PRC2-mediated H3K27me3 in ventral cells resulted in significant recovery in *Isl1* mRNA transcription [Fig. 2G] as well as reactivation of the *Isl1-nLacZ* reporter [Fig. 2H-I], respectively.

**Figure 2.**
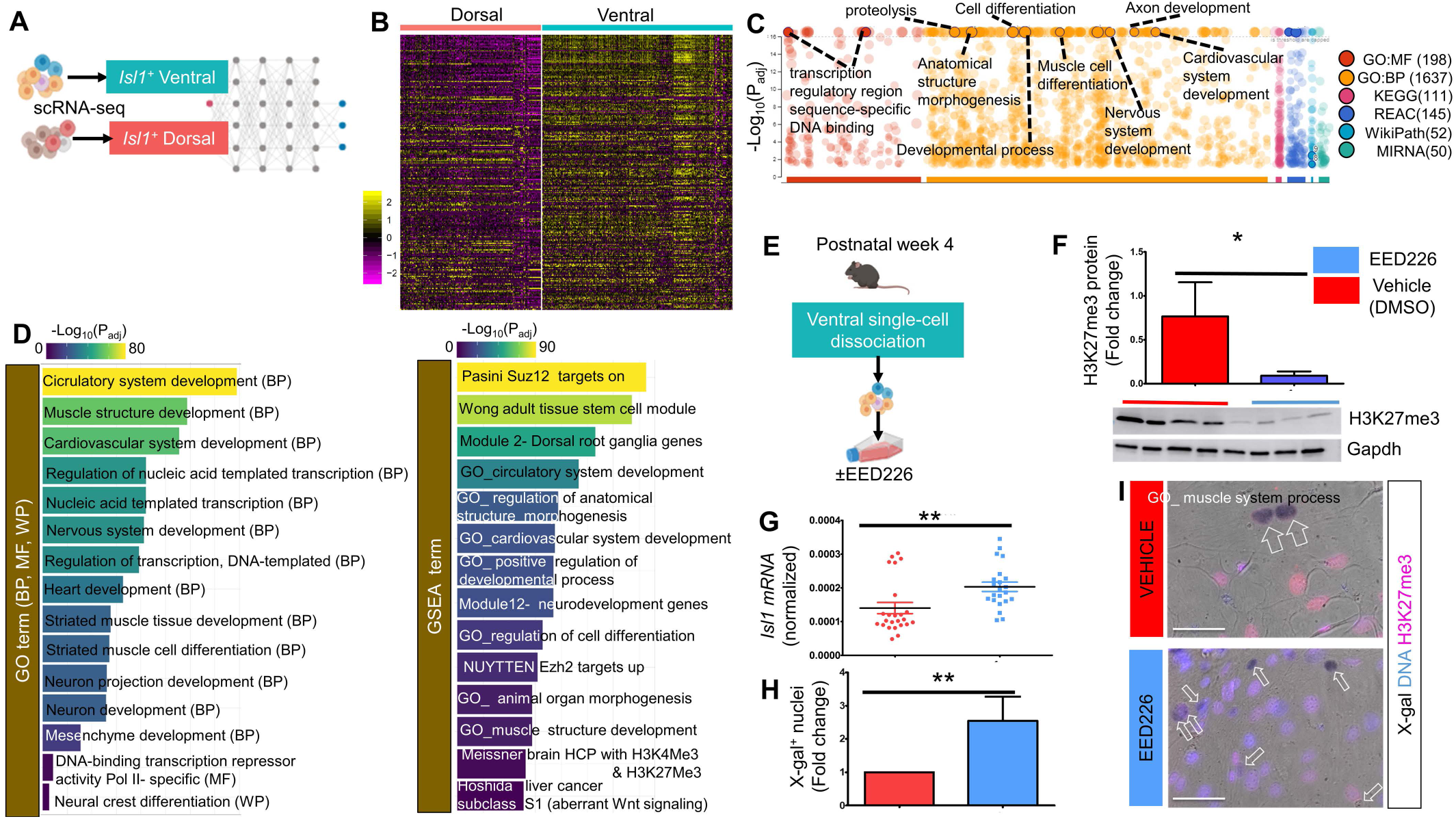
PRC2-mediated transcriptional silencing of ventral postnatal *Isl1* expression. A, Experimental outline. Dorsal vs ventral *Isl1^+^* cells were subselected from their respective scRNAseq datasets. **B**, Heatmap of the top 250 differentially expressed genes (DEGs, Wilcoxon rank sum test) between the dorsal and ventral *Isl1*^+^ cells (p-value adjustment was performed using Bonferroni correction based on the total number of genes in the dataset). **C**, Functional profiling of DEGs (false discovery rate adjusted *p*<0.05), visualized in a Manhattan-like plot. Functional terms are grouped and color-coded by data sources (total # terms/source shown in parenthesis). The g:SCS algorithm was applied for multiple testing correction of the adjusted enrichment p-values shown here in negative log10 scale [-log10(Padj)]. Values are capped at −log10(Padj)≤16. Examples of some of the most significant terms are noted. **D**, Bar graphs of over-represented Gene Ontology (GO, left) and gene-set enrichment analysis (GSEA, right) terms. The scale of −log10(Padj) values is numeric and color-coded. **E,** Illustration of the experimental outline to inhibit PRC2 activity in ventral postnatal *Isl1^+^* cells. **F,** Western blots of Gapdh and H3K27me3 after 5 days treatment with DMSO or 10μM EED226. **G-H**, Quantification of *Isl1* mRNA (F) and x-gal^+^ nuclei in each group. **I**, x-gal and H3K27me3 staining in each group. F-I, n=24 samples collected from 4 mice (12 samples/group). For qPCR, a technical replicate was included in each run. Scale bars, 150µm.

Therefore, based on the scRNAseq and functional experimental results we conclude that the Isl1-CPC-GRNs in the arterial pole/ ventral domain are transcriptionally silenced during postnatal growth, through the PRC2-mediated repression of the *Isl1* gene

Intriguingly, analysis of ChIP-seq datasets from whole embryonic and postnatal *Isl1^+/+^* mouse hearts showed that accumulation of the repressive histone mark H3K27me3 is evident as early as E10.5, and is selectively enriched at the *Isl1* promoter and heart field-specific enhancer regions, whereas the neural-specific, more distal regulatory elements of the gene are not repressed [Fig. S6]. This is in agreement with previous genetic screens for *Isl1* enhancers in transgenic reporter mice, showing permanent silencing of the heart field-specific *Isl1* enhancers after early cardiogenesis (Kappen and Salbaum, 2009). Therefore, based on these observations, it is conceivable that *Isl1-nLacZ* activity is sustained in both the ventral and dorsal postnatal cardiac domains through non-heart field regulatory elements. Moreover, given that ventral *Isl1-nLacZ* cells are a mixture of aSHF, CNC and venous pole derivatives [Figs. S2-3], the above findings indicate that silencing of postnatal *Isl1-nLacZ* is an arterial region-specific, rather than lineage- or cell type-specific, effect.

### Ventral, but not dorsal, Isl1 is translationally silenced after early cardiogenesis

We further observed that silencing of the heart field/ ventral *Isl1* regulatory elements is accompanied by complete loss of ventral Isl1 immunoreactivity, suggesting that the arterial pole *Isl1* domain is further regulated at the post-transcriptional level [Figs. 3A-B]. Because previous studies have suggested that Isl1 immunoreactivity differs between mouse strains (Khattar et al., 2011), we verified that loss of arterial but not venous pole Isl1 expression is conserved in human heart development [Fig. S7].

**Figure 3.**
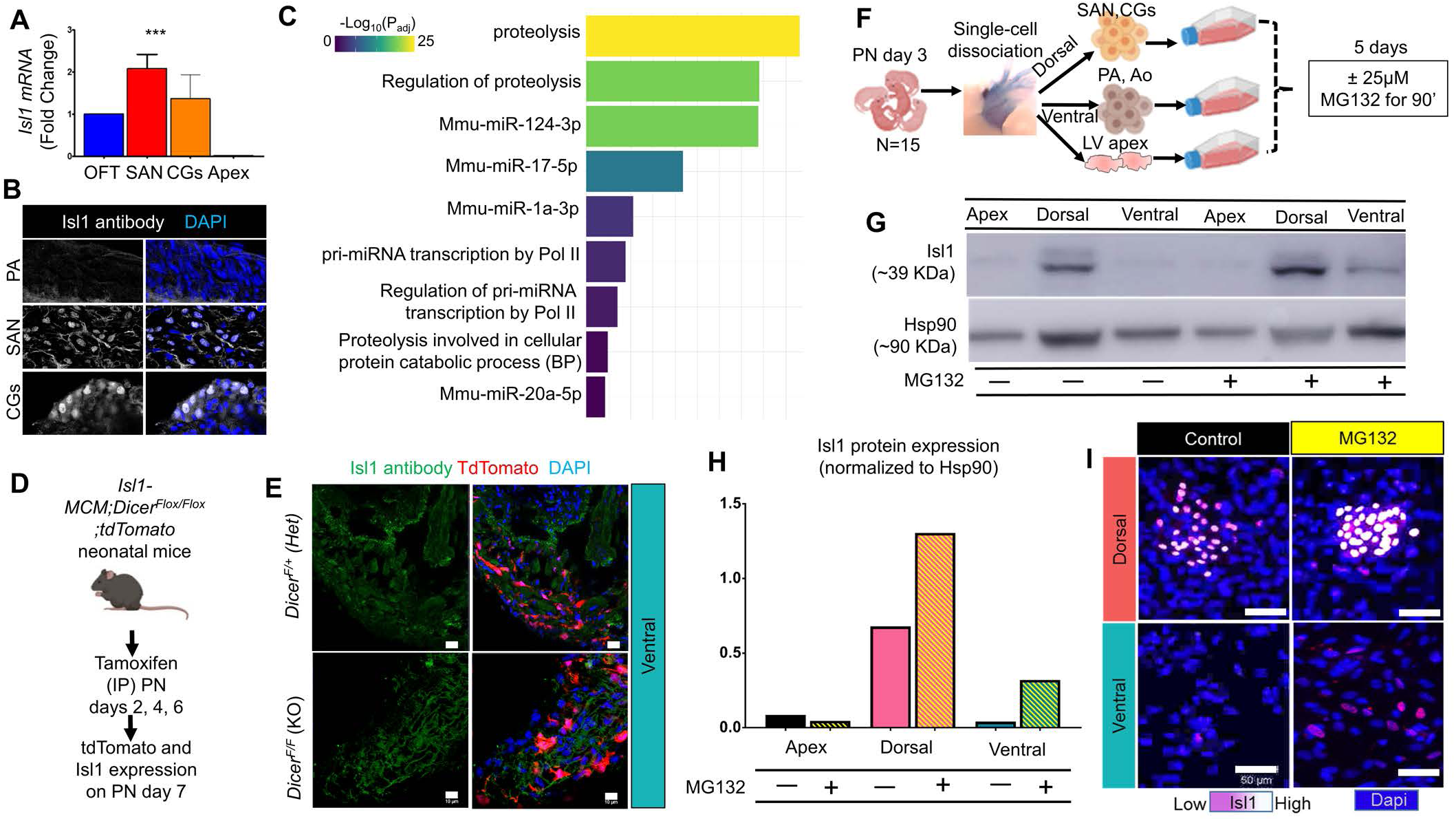
Proteasomal degradation of ventral, but not dorsal, *Isl1* expression. A-B, Ventral vs Dorsal Isl1 mRNA (A) and protein (B) expression (n=5 mice/group). **C**, Bar graphs of over-represented Gene Ontology (GO) terms. The scale of −log10(Padj) values is numeric and color-coded. **D-E**, Experimental design (D) and representative immunohistochemical analysis of Isl1 in *Isl1-MCM;Dicer^Flox/Flox^;tdTomato* mice (n= 4 heterozygous; n=8 Dicer-KO littermates). **F**, Illustration of the experimental outline to inhibit the ubiquitin-proteasome pathway in postnatal Isl1^+^ domains. **G-H**, Western blot analysis of Isl1 and Hsp90. **I**, Immunofluorescent analysis of Isl1. n= pool of 15 mice/group. Scale bars, 10µm (E); 50µm (I).

Functional enrichment analysis of DV *Isl1^+^* scRNAseq DEGs identified multiple miRNA-related GO terms, including miR-1a (Zhao et al., 2007) and miRNA-17 (Wang et al., 2010) [Fig. 3C; Table S3] which have been previously shown to play important roles in cardiac development. Importantly, miRNA-17, which is part of the miRNA 17∼92 cluster, has been shown to directly silence *Isl1* during aSHF development (Wang et al., 2010). Because miRNA biogenesis, including miR-1a and miRNA 17∼92, is largely dependent on the RNAse III nuclease *Dicer* (Du et al., 2015; Zhao et al., 2007), we conditionally knocked-out (cKO) *Dicer* in postnatal *Isl1* cells in order to functionally test the potential involvement of miRNAs in post-transcriptional silencing of ventral Isl1 expression. To do so, mice carrying a tamoxifen-inducible *Isl1-MerCreMer* (Laugwitz et al., 2005; Sun et al., 2007), a floxed *Dicer (Harfe et al., 2005)* and a *tdTomato Cre* reporter (Madisen et al., 2010) alleles (*Isl1-MCM;Dicer^Flox/Flox^;tdTomato)*] were generated and induced with intraperitoneal tamoxifen injections on PN 2, 4 and 6 [Fig. 3D. Excision of one or both *Dicer^Flox/Flox^* alleles was confirmed by tdTomato expression, as well as via RT-PCR amplification of the wild-type (WT), recombined and non-recombined floxed alleles. Confocal analysis on PN7 showed comparable ventral and dorsal tdTomato labeling between *Isl1-MCM;Dicer^Flox/Flox^;tdTomato* and *Isl1-MCM;Dicer^Flox/+^;tdTomato* hearts. However, loss of *Dicer* was not sufficient to reconstitute ventral Isl1 immunoreactivity [Fig. 3E]. Thus, silencing of ventral Isl1 protein expression in the postnatal heart is not mediated by *Dicer-* processed miRNAs.

Additionally to miRNAs, the DV *Isl1^+^* scRNAseq DEGs are enriched in multiple processes related to proteolysis [Fig. 2C; 3C; Table S3]. Notably, regulation of Isl1 protein stability and degradation has been shown to be essential for the establishment and differentiation of the aSHF in mouse embryos (Caputo et al., 2015). Accordingly, cells from the left ventricular apex (negative control for Isl1), dorsal (positive control for Isl1) and ventral domains of PN3 *Isl1^+/+^* hearts were isolated and treated for 90 minutes with 25μM of the ubiquitin-proteasome inhibitor MG132 (Clift et al., 2017), or vehicle (DMSO) [Fig. 3F]. Remarkably, Western blot [Fig. 3G, H] and immunocytochemical analyses [Fig. 3I] demonstrated that MG132-mediated inhibition of proteasomal degradation was sufficient to reconstitute ventral Isl1 protein expression, while at the same time upregulating Isl1 expression in dorsal cells [Fig. 3G-I].

Therefore, based on these findings we conclude that, in addition to their transcriptional silencing, the Isl1-CPC-GRNs in the arterial pole/ ventral domain are post-translationally silenced during fetal and postnatal growth, through the ubiquitin-proteasome pathway.

### *Isl1*^+^ CNCs contribute septal and trabecular cardiomyocytes that undergo proliferative expansion before, but not after, birth

Previous genetic studies in mice indicate that some postnatal *Isl1*^+^ cells are undifferentiated aSHF CPCs (Laugwitz et al., 2005). Others however, provide evidence against it (Li et al., 2019). To address this controversy, *Isl1-MCM mice* were crossed to the *tdTomato* or the dual-fluorescent *IRG* (Hatzistergos et al., 2015) reporter mice and induced with a single dose of tamoxifen on PN2 [Fig. S8A-B]. Analysis 48h later showed that, compared to the *Isl1-nLacZ* [Fig. S1D-E], *Isl1-MCM* recombination was partial because it selectively labeled the dorsal, but not ventral, Isl1^+^ cells [Fig. S8A-B]. This suggests that postnatal ventral *Isl1-MCM* expression is weaker than dorsal. Therefore, to capture the ventral *Isl1^+^* cells, we repeatedly administered tamoxifen on PN2 and PN4 and extended the period of fate-mapping to 2 weeks. Following this approach, we achieved comparable *Isl1-MCM* recombination in both domains [Fig. S8C-D]. However, compared to *Isl1-nLacZ, Isl1-MCM* hearts exhibited widespread labeling in CNC-derived TH^+^ neurons [Fig. S8 E-I], suggesting that postnatal Isl1-CPC-GRN activity is strongly associated to the postnatal growth of the CNC-derived sympathetic neuronal network. More importantly, compared to the *Isl1-nLacZ* which only labels CMs within the OFT base myocardium [Figs. S3A-BE; S8J], *Isl1-MCM* fate-mapping produced rare, isolated, tdTomato^+^ ventricular CMs, scattered within the compact and trabecular LV, RV and IVS myocardium [Fig. S8K-M]. These findings support previous reports that sustained postnatal Isl1-CPC-GRN activity is associated with sporadic CMs differentiation (Laugwitz et al., 2005).

Interestingly, *Isl1-MCM* CMs were occasionally found adjacent to *Isl1-MCM* TH^+^ nerves [Fig. S8J-M; movie S1]. This observation, along with the finding that a subset of *Isl1-nLacZ^+^* OFT base CMs likely originate from the CNC [Figs. S2-S3], led us to challenge the existing interpretations (Laugwitz et al., 2005) and hypothesized that that the *Isl1-MCM*-derived CMs reflect CNC-rather than aSHF-Isl1-CPC-GRN activity. Contributions of CNCs to the myocardial lineage have been previously shown in fish (Abdul-Wajid et al., 2018; Li et al., 2003; Tang et al., 2019) but remains a topic of debate in mice (Hatzistergos et al., 2018; Hatzistergos et al., 2016; Hatzistergos et al., 2015; Tang et al., 2019; Tomita et al., 2005).

Accordingly, to test this possibility, we first conducted clonal analysis of *Isl1*^+^ cardiac cells and their derivatives by crossing the *Isl1-Cre (Yang et al., 2006)* to the multicolor Cre-reporter mouse line *Confetti (Snippert et al., 2010)* [Fig. 4A]. Analysis of PN1 *Isl1-Cre;Confetti* hearts showed extensive labeling of the compact and trabecular RV myocardium in the form of multiple, single-colored CM clusters [Fig. 4B-C], consistent with previous findings that the RV myocardium develops from a relatively limited number of *Isl1^+^* aSHF CPCs which expand clonally before and after CM differentiation (Cai et al., 2003; Meilhac and Buckingham, 2018). However, some segments of the RV were not labeled by the *Confetti* [Fig. 4B-C]. Moreover, compared to *Isl1-Cre;tdTomato* (data not shown) and *Isl1-MCM;tdTomato* mice [Fig. S8C-I], fluorescent labeling of TH^+^ nerves in *Isl1-Cre;Confetti* mice was minimal in the RV, and absent in the LV and atria [Fig. 4B-C]. This suggests that *Isl1-Cre* expression is not sufficiently strong to excise the *lox-STOP-lox* cassette from the *Confetti*, thereby resulting in no fluorescent reporter labeling in some of the *Cre*-expressing cells (Snippert et al., 2010). Nonetheless, immunohistochemical analysis revealed single-colored, transmural RV clones within the *Isl1-Cre;Confetti* hearts which were heterogeneous clusters of α-sarcomeric actinin^+^ CMs and TH^+^ nerves [Fig. 4D-E]. Furthermore, to overcome the labeling limitations of the *Isl1-Cre;Confetti*, we repeated the clonal analysis by replacing the *Isl1-Cre* with the *Wnt1-Cre2* [Fig. 4A]. Compared to *Isl1-Cre, Wnt1-Cre2;Confetti* mice exhibited efficient labeling in CNC derivatives, including TH^+^ nerves [Fig. 4F]. Notably, most TH^+^ nerves were labeled as large, single-colored branched networks, indicating their clonal growth from a small number of CNCs (not shown). Analysis of PN7 *Wnt1-Cre2;Confetti* hearts revealed a small number of large, coherent, single-colored, myocardial clones within the RV, LV and IVS myocardium and trabeculae which were composed of Nkx2-5^+^/ α-sarcomeric actinin^+^ CM clusters and TH^+^ nerves [Fig. 4F-M].. Importantly, these large, transmural clones were wedge-shaped, with a wide epicardial and a narrower endocardial side, suggesting that each clone has developed from a single fetal CNC cell that underwent trabecular CM differentiation and then expanded proliferatively during ventricular compaction and septation. This finding is reminiscent of previous retrospective clonal analyses indicating two phases of clonal growth in the developing mouse heart and the presence of biventricular myocardial clones which do not derive from *Mesp1*-expressing mesodermal CPCs (Meilhac and Buckingham, 2018; Meilhac et al., 2003).

**Figure 4.**
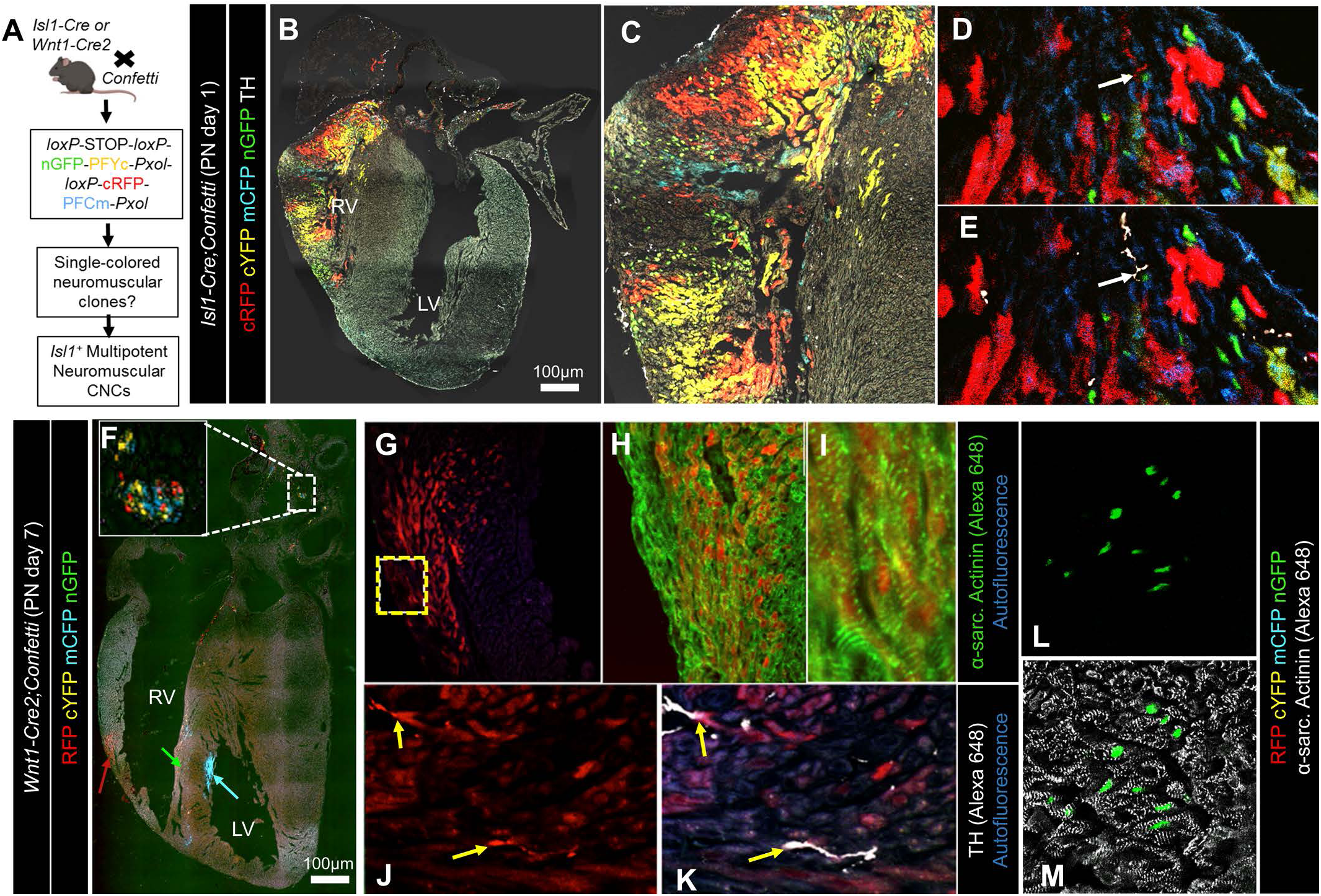
Isl1^+^ CNCs generate trabecular and septal cardiomyocytes with pre-but not post-natal, proliferative expansion capacity. **A,** Experimental outline to clonally track Isl1^+^ CNCs under the *Confetti*, *Isl1-Cre* (n=7) and *Wnt1-Cre2* (n=35) alleles. **B-C**, A postnatal day (PN)-1 *Isl1-Cre;Confetti* heart immunostained for tyrosine hydroxylase (TH). Expression of reporter genes is restricted in the right ventricle (RV), and indicates proliferative expansion of compact and trabecular RV cardiomyocyte clones. However, recombination is unsuccessful in most cardiac TH^+^ neurons inside and outside the RV. **D-E**, A red clone comprised of RV cardiomyocytes and a TH^+^ neural derivative, indicating common origin of the 2 cell types. **F-M**, A PN7 *Wnt1-Cre2;Confetti* heart immunostained for tyrosine hydroxylase (TH) or a-sarcomeric actinin. Inset on F depicts CNC-derived clones within a cardiac ganglion. **G-I**, A higher magnification of the wedge-shaped, transmural, coherent clone of a-sarcomeric actinin^+^ CNC-CMs (red arrow in F) indicating their clonal expansion from a single CNC progenitor. **J-K**, A higher magnification of the boxed area in G, indicating the clonal origin of CNC-CMs and TH^+^ neurons. **L-M**, A green clone of interventricular septum cardiomyocytes (green arrow in F) indicating their clonal expansion from a single CNC progenitor. Scale bars, 100μm.

Thus, based on the *Isl1-MCM* fate-mapping and *Confetti* clonal analyses experiments, we conclude that Isl1-CPC-GRN activity is sustained in the postnatal heart through the CNC, rather than the aSHF lineage; and that the Isl1-CNC-GRN endows the fetal and postnatal heart with limited biventricular cardiomyocyte and sympathetic nerve differentiation capacity. More importantly, activation of the Isl1-CNC-GRN during fetal growth and partitioning is coupled to CM proliferation and neural cell-cycle exit GRNs; whereas, sustained Isl1-CNC-GRN in the postnatal heart is coupled to sympathetic nerve proliferation and CM cell-cycle exit GRNs.

### Reconstruction of postnatal Isl1^+^ CNCs differentiation trajectory through scRNAseq

To challenge the interpretation of the lineage-tracing experiments that postnatal Isl1^+^ CNCs contribute to the Nkx2-5^+^ CM lineage, the *Isl1^+^* and *Nkx2-5^+^* fractions were sub-selected from the scRNAseq datasets, and subjected to pseudotime trajectory reconstruction through unsupervised, machine-learning approaches (Qiu et al., [Fig. 5A]. Overall, there were 442 *Isl1^+^*, 385 *Nkx2-5^+^* and 41 *Isl1^+^/Nkx2-5^+^* scRNAseq profiles which, when computed in 10 dimensions, were constructed into a principal graph with 3 branch points leading to 6 distinct cell states [Figs. 5B-C]. Branch point 1 diverges into cell states (CS)2, CS3 and CS4. Branch point 2 leads to CS5 and to branch point 3, which subsequently bifurcates into CS1 and CS6 [Fig. 5B]. Each state represents transcriptionally similar scRNAseq profiles, whereas the length of its trajectory projects the amount of transcriptional change as progress in pseudotime (Qiu et al., 2017a). Cluster-based analysis delineated that the 868 cells of the *Isl1^+^/Nkx2-5^+^* trajectory represent 14/15 of the UMAP clusters, with the macrophages (cluster 13) as the only exception [Fig. 5D, S9A]. The myocardial clusters (# 10 and 14) are ordered at the end of the CS4 trajectory [Fig. 5D, S9A], and therefore CS4 is classified as the myocardial lineage trajectory. CS4 branches out of branch-point 1, which is the point at which CS3 bifurcates into the myocardial and non-myocardial (CS1, 2, 5 and 6) trajectories [Fig. 5B]. Therefore, we classified CS3 as the root state.

**Figure 5.**
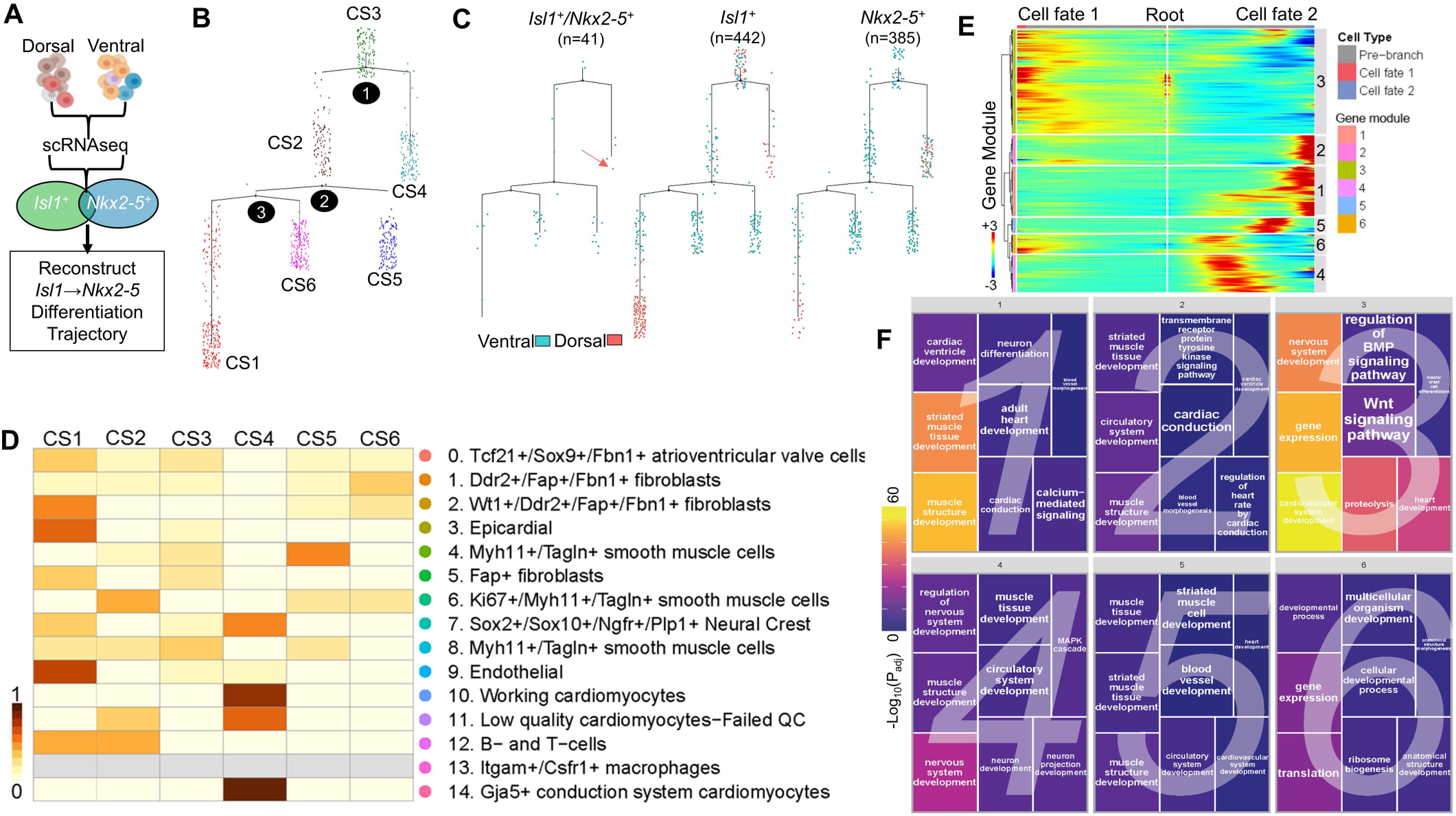
Reconstruction of postnatal *Isl1^+^* and *Nkx2-5^+^* pseudotime differentiation trajectories. **A,** Experimental outline. **B,** Ordering of the *Isl1^+^* and *Nkx2-5^+^* scRNAseq subsets (n=868 cells) on the reconstructed trajectory. The trajectory has 3 branch points (black circles) leading into 6 different cell states (CS, color-coded). CS3 is classified as the root. CS4 is classified as the myocardial state. CS1,2,5 and 6 are non-myocardial. **C**, Ordering of the *Isl1^+^*, *Nkx2-5^+^* and *Isl1^+^*/*Nkx2-5^+^* subsets, based on their dorsoventral identities. Overlap between *Isl1* and *Nkx2-5* expression is minimal (41/868 cells). In addition, 40/41 *Isl1^+^*/*Nkx2-5^+^* cells are of ventral origin indicating Isl1 protein degradation. Arrow points to a single *Isl1^+^*/*Nkx2-5^+^* dorsal cell at the tip of the myocardial trajectory. **D,** Heatmap of the distribution of UMAP clusters across each cell state. The myocardial trajectory (CS4) is composed by clusters 7, 9, 10, 11 and 14. Cluster 13 (grey) did not contribute to the trajectory. **E**, Multiway heatmap of branch point 1-dependent differentially expressed genes (false discovery rate-adjusted p<0.01), hierarchically clustered into 6 gene-modules. Gene modules 1 and 2 corresponds to the myocardial trajectory (CS4); gene module 3, to the root (CS3); and modules 4-6 to non-myocardial trajectories. **F**, Gene ontology term enrichment analysis of biological processes per gene module (false discovery rate adjusted p<0.01), visualized in a geometric treemap plot. Numbers 1-6 indicate gene modules. The scale of −log10(Padj) values is both geometrically and color-coded. Branched trajectories are plotted as a two-dimensional tree layout.

Interestingly, expression of *Isl1* and *Nkx2-5* showed minimal overlap within the reconstructed trajectories. In particular, only 41/868 cells co-expressed the two genes [Fig. 5C]. Of these, 4 were in the myocardial trajectory; 2 at the root cell state; and 35 in non-myocardial trajectories [Fig. 5C]. Moreover, 40/41 of the *Isl1^+^*/*Nkx2-5^+^* cells were of ventral identity [Fig. 5C] which, as demonstrated earlier (i.e. fig. 3), indicates that Isl1 is translationally silenced in these cells. These findings are reminiscent of previous developmental studies suggesting that the Isl1 and Nkx2-5 CPC-GRNs suppress each other during early CM differentiation of aSHF CPCs (Jia et al., 2018; Prall et al., 2007). Of note, consistent with our previous findings, a small fraction of *Isl1^+^* cells (27/483) and *Nkx2-5^+^* cells (5/426) co-expressed the proto-oncogene *Kit* (Hatzistergos et al., 2016; Hatzistergos et al., 2015).

Remarkably, from the 14 UMAP clusters in the *Isl1^+^*/*Nkx2-5^+^* pseudotime trajectories, only five clusters-#7 (CNCs), #9 (venous endothelial), #10 (working CMs); CMs); #11 (low-quality/dying CMs) and #14 (Gja5^+^ conduction system CMs)-follow the the myocardial lineage trajectory (CS4) [Figs. 5D; S9A-B]. Specifically, the myocardial cells (clusters 10 and 14, as well as the low-quality cluster 11) are ordered at the end of the trajectory, consistent with their terminal differentiation state [Fig. S9A-B]. Whereas, *Isl1*^+^ CNCs are placed at the start of the trajectory resembling a progenitor state [Fig. S9A-B]. Importantly, all except one *Isl1^+^* CNC profiles are of venous pole/ dorsal identity [Fig. S9C], indicating functional Isl1 protein expression. Similarly, the single *Isl1^+^* cell from the endothelial cluster that is placed in the myocardial trajectory is also of venous pole/ dorsal identity [Fig. S9C], thereby excluding an arterial pole/ aSHF CPC ancestry. Nonetheless, to investigate further if any of the postnatal Isl1^+^ cells represent aSHF-CPCs, we also analyzed the *Isl1^+^/Nkx2-5^+^* trajectory on the basis of *Kdr* and *Pdgfra* expression. These 2 receptors are transiently co-expressed on the surface of the majority of multipotent cardiogenic mesoderm CPCs, including aSHF, during mouse and human development (Kattman et al., 2011; Prall et al., 2007; Zhang et al., 2016). Overall, there were 415 *Pdgfra^+^*, 10 *Kdr^+^* and 4 *Kdr^+^/Pdgfra^+^* cells [Fig. S9D]. However, the *Kdr^+^/Pdgfra^+^* fraction represent a venous pole/dorsal cell subset which belong to UMAP clusters 1, 3, 9 and are ordered at the end of the non-myocardial trajectory CS1 [Fig. S9D].

Thus, based on the above lineage-tracing and scRNAseq studies, we conclude that a subset of venous pole/ dorsal *Isl1^+^* CNC-derived cells in the embryonic and postnatal heart are endowed with a multipotent neuromuscular progenitor cell state. In contrast, we could not find any evidence supporting the presence of arterial pole/ aSHF-derived postnatal CPCs.

### Disruption of Wnt/β-catenin-Isl1 feedback loop promotes CNC cardiomyocyte differentiation

To identify potential molecular mechanisms controlling the fate of postnatal *Isl1^+^* CPCs, we performed DEG analysis (Qiu et al., 2017a) on branch-point 1, where the root state bifurcates toward myocardial or non-myocardial trajectories [Figs. 5B; S9A]. There were 1,924 branch-dependent DEGs (q-value <0.01) which were hierarchically clustered on a multiway heatmap into 6 gene-modules. Each gene module indicates similar pseudotime trajectory-dependent expression patterns [Fig. 5E; Table S5]. Gene modules 1 and 2 exhibit upregulation in cell fate 2 and downregulation in cell fate 1 genes. Gene module 3 exhibits upregulation in root state (center) and cell fate 1 pre-branch genes, but downregulation in cell fate 2 genes. Whereas, gene modules 4, 5 and 6 exhibit upregulation in cell fate 2 pre-branch genes [Fig. 5E]. Functional enrichment analysis indicated that gene modules 1 and 2 are enriched in biological processes related to the myocardial lineage, including embryonic and adult heart development; heart contraction, cardiac conduction and cardiac ventricle morphogenesis [Fig. 5F, Table S6]. Gene modules 4, 5 and 6 are more heterogeneous as indicated by their enrichment in pre-branch genes related to myocardial, neuronal and vascular biological processes [Fig. 5F, Table S6]. Whereas, gene module 3 is enriched in biological processes related to the cardiac and neural crest differentiation and proliferation, as well as proteolysis, gene regulation, Wnt and BMP signaling pathways [Fig. 5F, Table S6]. Thus, based on the above pseudotime trajectory-dependent expression profiles and functional enrichment analyses, we conclude that gene module 3 captures the root state gene-signature underlying the decisions of postnatal *Isl1^+^* cells toward myocardial or non-myocardial fates; gene modules 1 and 2 capture the terminally differentiated CM states; and gene modules 4-6 capture the remaining cell states.

Based on scRNAseq functional enrichment analysis, the root state (gene module 3) of the *Isl1^+^/Nkx2-5^+^* trajectory receives input from the Wnt and BMP signaling pathways [Fig. 5F, Table S6]. During normal heart development, canonical Wnt signaling directly activates *Isl1* to inhibit CM differentiation and promote the expansion of aSHF CPCs (Lin et al., 2007), whereas repression of Wnt by canonical BMP signaling is required for downregulation of *Isl1* and induction of cardiomyogenesis (Jain et al., 2015). Therefore, we sought to functionally test the role of canonical Wnt signaling in the fate of *Isl1^+^* CNC-derived neuromuscular precursors. First, we genetically fate-mapped CNCs in the presence or absence of a Wnt/β-catenin gain-of-function mutation, introduced through a previously described *Wnt1/GAL4/Cre-11* transgene (Lewis et al., 2013). Consistent with the *Confetti*-and scRNAseq-based fate-maps, CNCs under normal Wnt signaling contributed to all expected derivatives in the OFT and cardiac autonomic nervous system, as well as to Nkx2-5^+^ CMs within the trabeculated [Fig. 6A] and compact LV, RV and IVS myocardium [Fig. 6B]. However, ectopic activation of *Wnt* produced an ∼147-fold reduction in CMs differentiation while OFT and neuronal contributions were unaffected [Fig.6C-D] (0.07±0.04% vs. 9.76±0.77% myocardial labeling under the *Wnt1/GAL4/Cre-11* and *Wnt1-Cre2* drivers, respectively; *p*<0.0001).

**Figure 6.**
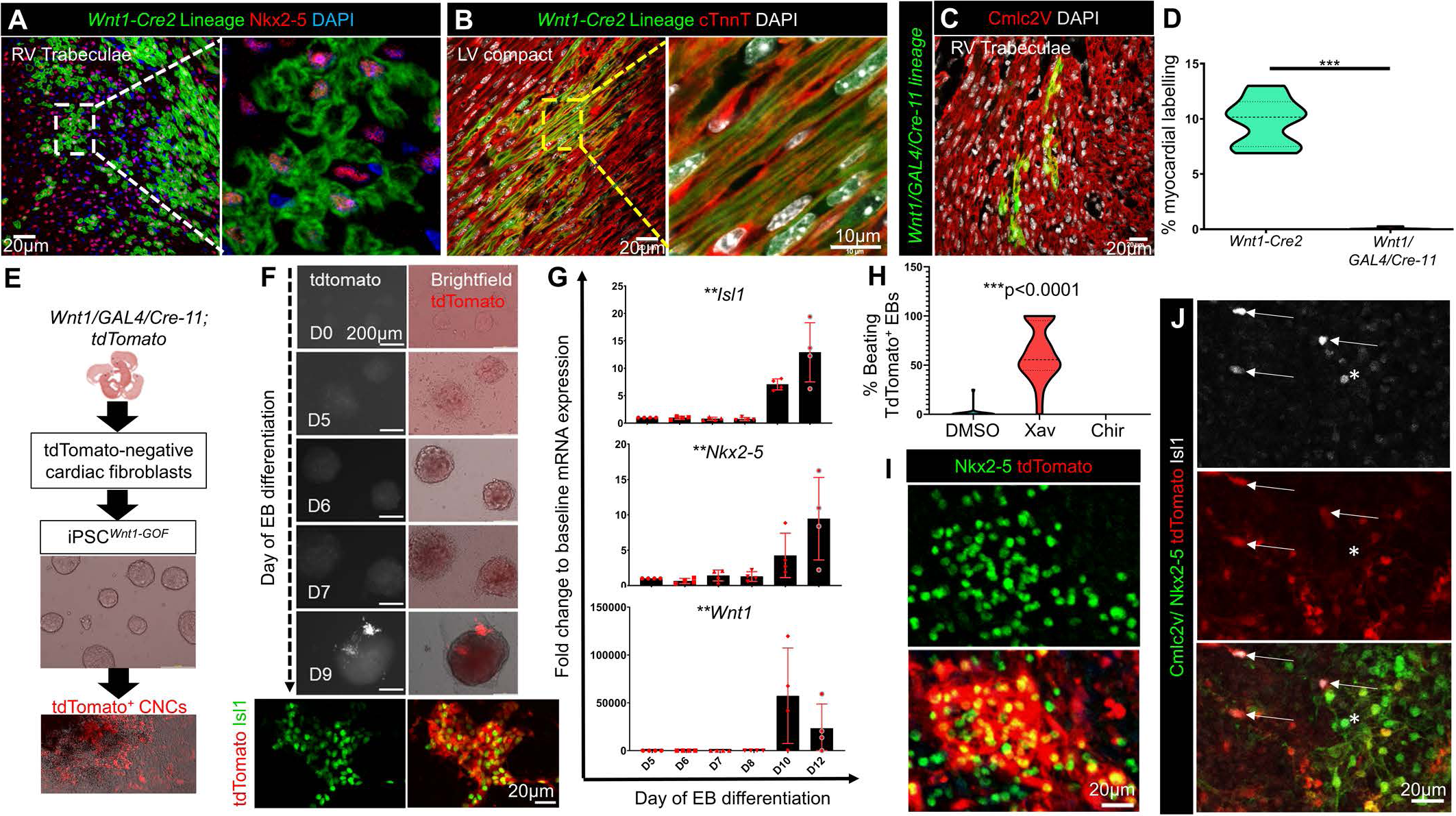
Cardiomyocyte differentiation of Isl1^+^ CNCs is controlled by Wnt/β-catenin. **A-B,** CNCs contribute to the normal development of trabecular and compact ventricular myocardium, as indicated by expression of the Cre reporter in clusters of Nkx2-5^+^ (A) and cardiac troponin T^+^ (B, cTnnT) cardiomyocytes (CMs), in a postnatal day (PN) 7 *Wnt1-Cre2;TdTomato* heart. **C**, CNC-CM differentiation is diminished in response to ectopic activation of Wnt/β-catenin in CNCs through a *Wnt1/Gal4/Cre-11* transgene. **D**, Violin plot of tdTomato^+^ myocardium as a percentage of the total area of the heart, between *Wnt1-Cre2* and *Wnt1/GAL4/Cre-11* reporter neonates (n=3 mice/group, 2-3 slides/heart, \*\*\**p*= 0.0005, Mann-Whitney test). **E**, Outline of iPSC*^Wnt1-GOF^* modeling experiments. **F**, Time-course of iPSC-derived embryoid bodies (EB) differentiation into tdTomato^+^/Isl1^+^ CNCs. **G**, Temporal analysis of *Isl1*, *Nkx2-5* and *Wnt1* expression during EB differentiation (n=4/time point, Kruskal-Wallis, \*\**p*<0.005). **H**, Violin plot of % beating EBs containing tdTomato^+^ cells in response to DMSO, Xav-939 (Xav) or Chir99021 (Chir) (n= 143, 129 and 55 EBs, respectively, Kruskal-Wallis, \*\*\**p*<0.0001). **I**, Expression of Nkx2-5 in tdTomato^+^ derivatives following treatment with XAV-939. **J**, Silencing of Isl1 expression following cardiomyogenic differentiation of tdTomato^+^ cells. Arrows point to undifferentiated Isl1^+^/tdTomato^+^ cells (no expression of Nkx2-5 and cmlc2v). Asterisk indicates a tdTomato-negative/Isl1^+^//Cmlc2v^+^/Nkx2-5^+^ heart-field ventricular CPC. Values are mean±SEM.

To better explore the effects of Wnt signaling on CNC cardiomyogenesis, we generated *Wnt1/GAL4/Cre-11;tdTomato* induced pluripotent stem cells (iPSC*^Wnt1-GOF^*) and differentiated them into CNCs [Fig. 6E] (Hatzistergos et al., 2018; Hatzistergos et al., 2015). Specification of iPSC*^Wnt1-GOF^* into Isl1^+^ CNCs commences within 9 days of EB differentiation, as indicated by induction of tdTomato; activation of *Wnt1, Isl1* and *Nkx2-5* mRNAs; and co-localization of tdTomato with Isl1 [Fig. 6F-G]. By day 12, tdTomato is expressed in 61.81±5.76% EBs. Treatment of day 9 EBs with the GSK-3 inhibitor CHIR99021 or DMSO, did not promote cardiac differentiation, as indicated by the lack of spontaneously beating tdTomato^+^ cells [Fig. 6H-I; Movie S2]. In contrast, antagonism of Wnt with the tankyrase inhibitor XAV939 resulted in an ∼86-fold increase in the emergence of spontaneously beating tdTomato^+^/Nkx2-5^+^ cells (*p*<0.0001) [Fig. 6H-J; Movie S3]. Notably, XAV939-mediated induction of Nkx2-5 was accompanied by downregulation of Isl1 immunoreactivity [Fig. 6J], further supporting the scRNAseq *Isl1^+^/Nkx2-5^+^* trajectory that Isl1 expression is silenced in Nkx2-5^+^ cells [Figs 5B and S9B].

Finally, we conditionally deleted *Isl1* from CNCs (*Isl1-cKO*) by introducing an *Isl1^Fl/Fl^* allele (Sun et al., 2008) to *Wnt1-Cre2;tdTomato* and *Wnt1/GAL4/Cre-11;tdTomato* mice [Fig. 7A]. *Wnt1-Cre2;tdTomato;Isl1^Fl/+^* mutants are produced at the expected Mendelian ratios with grossly normal hearts; whereas, *Wnt1-Cre2;tdTomato;Isl1^Fl/Fl^* embryos succumb to death between E14.5-E15.5 [Fig. 7B]. Analysis at E14.5 showed that, compared to their heterozygous littermates, *Isl1-cKO* mutant hearts exhibit increased concentration of tdTomato^+^ CNCs in the AV cushions; and abnormal RV and LV trabecular networks, including a marked reduction in the contribution of tdTomato^+^ CNCs into Nkx2-5^+^ trabecular and IVS CMs as well as muscular IVS defects [Fig. 7C-D]. Remarkably, loss of *Isl1* under ectopic Wnt activity from the *Wnt1/GAL4/Cre-11* driver (*Isl1-cKO;Wnt1-GOF)*, results in normal Mendelian genotypic ratios [Fig. 7B], but *Isl1-cKO;Wnt1-GOF* hearts exhibit a near-complete loss of tdTomato^+^ sympathetic nerves and CMs, and die shortly after birth as previously reported (Sun et al., 2008). Interestingly however, in both *Wnt1-Cre2* and *Wnt1/GAL4/Cre-11* mice, heterozygous deletion of *Isl1* was accompanied by increased differentiation of CNCs into tdTomato^+^ myocardium compared to *Isl1*^+/+^ controls [Fig.7E-G], indicating that, similar to its function in other neural crest derivatives (Song et al., 2009), *Isl1* gene dosage influences the specification of CNCs toward CM vs non-CM fates.

**Figure 7.**
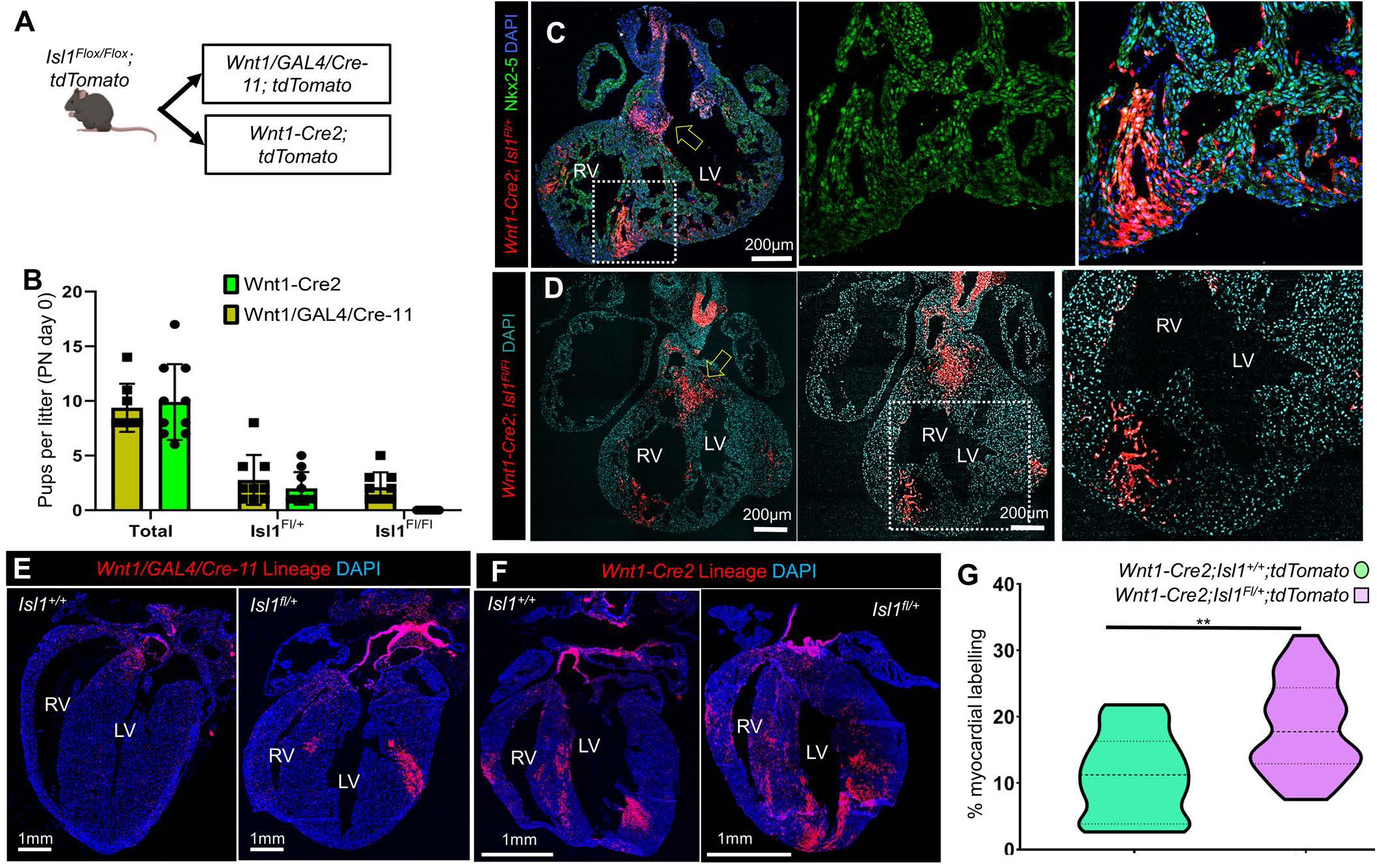
*Isl1* gene dosage regulates the cardiomyogenic fate of CNCs. A, Experimental outline. **B,** Pup survival per litter following conditional deletion of Isl1 in CNCs by *Wnt1-Cre2* (n=43) and *Wnt1/GAL4/Cre-11* (n=74). CNC-specific Isl1-cKO is developmental lethal, but is rescued by ectopic Wnt. **C-D**, Expression of tdTomato in E14.5 *Wnt1-Cre2;tdTomato;Isl1^Fl/Fl^* and *Wnt1-Cre2;tdTomato;Isl1^Fl/+^* hearts indicates Isl1-cKO is characterized by abnormal migration and differentiation of CNCs into trabecular and septal cardiomyocytes (CMs). **E-F**, Expression of tdTomato in *Isl1^+/+^* and *Isl1^Fl/+^* postnatal hearts under the *Wnt1-Cre2* and *Wnt1/GAL4/Cre-11*. Loss of myocardial derivatives by ectopic Wnt through *Wnt1/Gal4/Cre-11* is partially rescued when the *Isl1* gene dosage is reduced to half (E). Similarly, a reduction of Isl1 gene dosage by half promotes an increase in CNC-CM differentiation by 66% in *Wnt1-Cre2* mice (F). **G**, Violin plot of *Wnt1-Cre2* myocardial labeling as a percentage of the total area of the heart, between *Isl1^+/+^* and *Isl1^Fl/+^* (n=3 mice/group, 4-5 slides/heart, ***p= 0.0022, paired t-test). Values are mean±SEM.

Therefore, based on the above findings we conclude that the venous pole-derived/ dorsal Isl1^+^ CNCs play an important developmental role in myocardial trabeculation, compaction and sympathetic innervation, and suggest that dorsal Isl1^+^ CNCs are sustained long-term through a forward reinforcing loop between Isl1 and Wnt/β-catenin signaling, the disruption of which drives CMs differentiation.

## DISCUSSION

The main finding of our study is that organ-founding CPC-GRNs are sustained in the venous, but not arterial pole region of the postnatal heart, through CNC specific-but not the mesoderm-specific *Isl1* CPC-GRNs. This is an unexpected result given the existing interpretation that Isl1^+^ CPCs are aSHF-derived (Laugwitz et al., 2005; Meilhac and Buckingham, 2018); and that CM differentiation is thought as a transient, early developmental cardiomyogenic process (Li et al., 2019).

We focused on *Isl1* to gauge myocardial plasticity, because this gene is expressed in most, if not all, mammalian CPC-GRNs during development and remains active in subsets of incompletely characterized arterial and venous pole derivatives after birth (Laugwitz et al., 2005; Ma et al., 2008; Prall et al., 2007; Sun et al., 2007). Importantly, studies in mice have shown that each CPC lineage utilizes distinct *cis-* regulatory elements to drive *Isl1* expression in a spatially and temporally controlled manner. Heart field-specific enhancers are found ∼10kb to the 3’-end of the mouse *Isl1* gene and are highly conserved throughout evolution. They are originally activated in migrating lateral plate mesoderm cells (the precursors to both FHF and SHF), before becoming restricted to aSHF and pSHF CPCs (Kang et al., 2009; Kappen and Salbaum, 2009; Ma et al., 2008; Prall et al., 2007; Verzi et al., 2005). They are activated through GATA and forkhead transcription factor binding sites (Kang et al., 2009; Kappen and Salbaum, 2009), and negatively regulated indirectly by Nkx2-5, through incompletely understood mechanisms (Prall et al., 2007). Neural/CNC enhancers are found distally (∼0.5Mb) from the 3’-end. They are an evolutionary innovation to the *Isl1* gene, exapted from ancient transposable element non-coding sequences (Bejerano et al., 2006; Uemura et al., 2005). They are regulated through homeodomain response elements, with high affinity to Phox2b, Hox and Lhx and lower affinity to Nkx2-5 (Berger et al., 2008; Chen and Schwartz, 1995; Uemura et al., 2005). In addition, *Isl1* expression is regulated through a β-catenin/Lef1 responsive promoter proximal to its 5’ *ATG* transcription start site (Lin et al., 2007).

In keeping with our findings, previous studies provided evidence that silencing of heart field-specific Isl1 expression starts as early as E8.5 in mice. Particularly, aSHF enhancer-specific β-galactosidase reporters were shown to exhibit broader expression than the endogenous Isl1 protein at the looped heart stage. However, this discrepancy has been heretofore attributed to the longer β-galactosidase half-life (Kang et al., 2009; Zhuang et al., 2013). We now show that, in fact, it reflects the developmental activation of the ubiquitin-proteasome pathway, selectively in the arterial-but not venous-pole region, which leads to Isl1 protein degradation, in both mice and humans. A similar mechanism has been shown to regulate Isl1 protein levels in aSHF-CPCs at the crescent and linear heart tube stages (Caputo et al., 2015). Here, we further show through lineage-tracing experiments that arterial pole-specific Isl1 degradation affects SHF and CNC derivatives equally, indicating that this post-translational mechanism is specific to the arterial pole domain rather than the lineage or type of cells, and overrides *Isl1* expression from both its mesoderm and neural regulatory elements. Notably, expression of *Isl1* is also silenced through the miRNA 17∼92 cluster during aSHF-CPC differentiation (Wang et al., 2010). However, in our study, conditional deletion of *Dicer* was not sufficient to drive protein expression in ventral postnatal *Isl1^+^* cells, suggesting that this miRNA mechanism is restricted to aSHF enhancer– specific *Isl1* gene products during the early CM differentiation stages of aSHF-CPCs.

The permanent, developmental silencing of the arterial pole Isl1^+^ domain is further supported by previous studies showing that the activity of cardiac mesoderm-specific enhancers diminishes before E12.5 in mice (Kappen and Salbaum, 2009). However, at first glance, this finding seems somewhat contradictory to ours, since we show that developmental silencing of Isl1 is primarily post-translational rather than transcriptional. In fact, we show that *Isl1-nLacZ* and *Isl1* mRNAs continue to be abundantly expressed in both heart field (SHF/SAN) and CNC derivatives throughout development; and that transcriptional silencing of *Isl1* occurs after birth through a relatively slow process which takes months to complete and involves DV differences in postnatal PRC2 activity. We further show that postnatal *Isl1* silencing occurs in a regionalized fashion, selectively in the arterial-but not venous-pole domain and affects SHF and CNC derivatives equally. Interestingly however, a limitation of our study compared to Kappen *et al*. is that our findings are based on the *Isl1-nLacZ* knock-in which exhibits broad activity from both the neural-and mesoderm specific regulatory elements. Therefore, the seemingly discrepant findings may indicate that late *Isl1* expression is sustained in the developing and postnatal heart through non-aSHF regulatory elements. Support to this explanation comes from fate-mapping the postnatal *Isl1-nLacZ* activity through an *Isl1-MCM* version of the allele. These experiments demonstrated that postnatal *Isl1* activity is broadly associated with CNC-derived, TH^+^ sympathetic neurogenesis, although a remarkable, but less frequent, contribution to biventricular and IVS cardiomyogenesis is also made. More importantly, clonal analysis with the *Confetti* multicolor lineage tracer, as well as scRNAseq-based lineage trajectory reconstruction, indicated differentiation of TH^+^ neurons and CMs from a common Isl1^+^ CNC progenitor, demonstrating that neural/CNC enhancers activate both neurogenic and CM *Isl1-*CPC-GRNs.

Differentiation of CNCs to CMs has been previously shown in zebrafish (Abdul-Wajid et al., 2018; Li et al., 2003; Sande-Melón et al., 2019; Sato and Yost, 2003; Tang et al., 2019) and mice (Hatzistergos et al., 2015; Tamura et al., 2011; Tang et al., 2019; Tomita et al., 2005). Developmentally, CNCs primarily contribute trabecular CMs in both species; whereas in mice, they further contribute IVS CMs, a structure that has not evolved in the univentricular zebrafish heart (Abdul-Wajid et al., 2018; Hatzistergos et al., 2015; Tang et al., 2019). Consistently, we show that migration and CM differentiation are compromised in Isl1-cKO CNCs and that these mice die at midgestation with abnormalities in AV cushions, IVS and trabecular myocardium. Previous studies in mouse embryos have shown that IVS and trabecular CMs are clonally related and that these late-forming CMs develop progressively, during E10.5-PN7, as transmural, wedge-shaped clusters, with a wide epicardial and a narrower endocardial side (Meilhac et al., 2003). This description matches the CM clones produced from CNCs [e.g. Fig. 8]. Moreover, *Confetti* analysis illustrated that these are primarily single-colored coherent clones, which as previously suggested (Meilhac et al., 2003), indicates that upon initial differentiation, CNC-CMs undergo a phase of massive proliferative expansion [e.g. Fig. 4F]. Remarkably, this proliferative expansion of CNC-CMs has been shown to underlie adult heart regeneration in zebrafish (Abdul-Wajid et al., 2018; Sande-Melón et al., 2019), suggesting that it could also be targeted therapeutically to enhance the adult heart regenerative capabilities in mammals.

Interestingly, we found that mice with heterozygous loss of *Isl1* in CNCs develop normally, but exhibit a significant increase in the number of Nkx2-5^+^ CNC-CM clones [e.g. Fig. 8F-G]. This indicates that *Isl1* gene dosage influences the CM vs non-CM fates of CNCs and is reminiscent of the role of *Isl1* in specifying motor neuron vs V2a interneuron identities in neural crest cells (Song et al., 2009). Particularly, Song *et al*. showed that heterozygous loss of *Isl1* resulted in higher Lhx3 and Nkx6.1 levels which consequently altered the relative stoichiometries of the Ldb1/Lhx3/Isl1 complexes in favor of V2a interneurons vs motor neuron differentiation. Similarly, formation of Ldb1/Isl1 complexes were recently shown to regulate Isl1 protein levels by controlling its proteasomal degradation during aSHF-CPC differentiation (Caputo et al., 2015). Here, we were to determine through scRNAseq and pharmacologic experiments that a similar mechanism contributes to the sustained expression of Isl1 in dorsal CNCs. Finally, as predicted through scRNAseq-based lineage trajectory reconstruction, we further show that the CNC-CM fate is additionally regulated through a forward reinforcing loop between Wnt/β-catenin and Isl1. In particular, mice with ectopic activation of Wnt in CNCs are viable and fertile but exhibit ∼147-fold decrease in CNC-CMs. Remarkably, impaired CM differentiation could be rescued by eliminating one of the *Isl1* alleles, as well as through pharmacologic inhibition of Wnt in iPSCs-derived Isl1^+^ CNCs. Of note, the grossly normal cardiovascular phenotype of Wnt1-GOF mice suggests that CNC-CMs are not essential for the development of trabeculated and IVS myocardium since it appears to be compensated through other CM sources. Interestingly, loss of CNC-CMs does not affect heart development in zebrafish either, but leads to adult onset hypertrophic cardiomyopathy (Abdul-Wajid et al., 2018) and impaired postnatal cardiac regenerative capacity (Sande-Melón et al., 2019; Tang et al., 2019). We have also observed signs of LV and IVS hypertrophy in Wnt1-GOF mice [e.g. Fig. 8E and unpublished observations], however whether this is normal or pathological was not examined in this study.

In summary, our findings provide important insights into the deployment and withdrawal dynamics of the organ-founding CPC-GRNs during the embryonic development and postnatal growth of the mouse heart; and demonstrate that the postnatal heart sustains a pool of organ-founding CNCs in the inflow region. These “dorsal CNCs” could be stimulated to proliferate massively after they differentiate by evoking compaction-like proliferative signals, thereby providing a potential regenerative medicine approach to the vulnerability of humans to heart disease.

## Acknowledgements

This work was supported by the National Institutes of Health grants R01 HL107110, R01 HL094849, R01 HL110737, R01 HL084275, 5UM HL113460; grants from the Starr Foundation and the Soffer Family Foundation (all awarded to J.M.H.). This work was further supported by R01 CA125970 (J.W.H.); a University of Miami Medical Scientist Training Program (M.A.D.), the Sheila and David Fuente Graduate Program in Cancer Biology (M.A.D.); a Center for Computational Science Fellowship (M.A.D.); and a generous gift from Dr. Mark J. Daily (J.W.H). The Sylvester Comprehensive Cancer Center also received funding from the National Cancer Institute Core Support grant P30 CA240139. The Bascom Palmer Eye Institute also received funding from NIH Core Grant P30EY014801 and a Research to Prevent Blindness Unrestricted Grant. We thank Dr Sylvia M. Evans from the Skaggs School of Pharmacy and Pharmaceutical Sciences, UCSD, La Jolla, California, USA; and Dr Chenleng Cai from the Icahn School of Medicine at Mount Sinai, New York, USA; for providing the *Isl1-nLacZ* mice. We thank the Mutant Mouse Regional Resource Centers (MMRRC) for the cryopreserved *Mef2c-AHF-Cre* material. We also thank the ENCODE Consortium and the ENCODE production laboratory of Dr Bing Ren at UCSD for generating the ChIP-seq datasets. We acknowledge the support of the Biostatistics & Bioinformatics and Oncogenomics Shared Resources at the Sylvester Comprehensive Cancer Center and the Center for Computational Science High Performance Computing Group at the University of Miami.

## Author Contributions

K.E.H conceived the project (with input from J.M.H.), performed experiments, analyzed and interpreted the data and wrote the manuscript. M.A.D. analyzed and interpreted the data and edited the manuscript. J.W.H. and J.M.H provided resources, interpreted the data, and wrote the manuscript.

## Methods

### Experimental Model and subject details

#### Mice

All animals were maintained in an AAALAC-approved animal facility at the University of Miami, Miller School of Medicine, and procedures were performed using IACUC-approved protocols according to NIH standards. The *Isl1-nLacZ* mice have been described before (Sun et al., 2007). The *IRG* (Stock #008705), *tdTomato* (Stock #007914)*, Confetti* (Stock #017492), *Wnt1-Cre2* (Stock #022501), *Isl1-MerCreMer* (Stock #029566), *Isl1^fl/fl^* (Stock #028501), *Dicer^fl/fl^* (Stock #006366), *Isl1-Cre* (Stock #024242) and *Wnt1/GAL4/Cre-11* (Stock #003829) mice were obtained from the Jackson Laboratory. The *Mef2c-AHF-Cre* mice (Verzi et al., 2005) were cryorecovered at the University of Miami, Sylvester Comprehensive Cancer Center, Transgenic animal facility, from material obtained from the Mutant Mouse Regional Resource Centers (MMRRC). The *Wnt1-Cre2* mice were bred through the female germline. The *Mef2c-AHF-Cre* mice were bred through the male germline. Genotyping was performed by an independent provider via an automated real-time PCR system (Transnetyx). All analyses were performed in age-matched males and female littermates from multiple litters.

#### Human Fetal Heart Tissue

Human fetal heart tissues (15-22 weeks of gestation) were obtained from authorized sources (Advanced Bioscience resources, Inc, Alameda, CA) following IRB approval. Upon arrival, tissues were fixed in 10% buffered formalin for ∼24h, embedded in paraffin, and cut into 4-5μm-thick sections as previously described (Hatzistergos et al., 2010).

### Method Details

All procedures were performed using IACUC-approved protocols according to NIH standards.

#### Tamoxifen Injection

For *Isl1-MerCreMer* recombination, 2 day old neonatal mice were injected subcutaneously with 50μl of tamoxifen (Sigma), dissolved in peanut oil (Sigma) at a concentration of 20mg/ml, as previously described(Hatzistergos et al., 2016). Animals were euthanized 48h later. For fate-mapping experiments, tamoxifen injections were repeated on days 2 and 4, and animals were euthanized 2 weeks later. For Dicer-Knockout experiments, injections were repeated on days 2, 4 and 6, and animals were euthanized on day 7.

#### Immunohistochemistry and immunocytochemistry

For Immunocytochemical analysis, cells were fixed in 4% paraformaldehyde, blocked for 1h with 10% normal donkey serum, and processed for immunostaining as described before (Hatzistergos et al., 2016; Hatzistergos et al., 2015). Immunofluorescence analysis of mouse heart tissues was performed in 10μm-thick cryosections as described before (Hatzistergos et al., 2016; Hatzistergos et al., 2015). Immunofluorescence analysis of human fetal heart samples was performed in 4-5μm-thick formalin-fixed, paraffin-embedded tissue sections as previously described (Hatzistergos et al., 2010; Hatzistergos et al., 2015). Antigen unmasking was performed by microwaving the slides for 2×10min in citrate buffer Solution, pH=6 (ThermoFisher). Sections were then blocked for 1h at RT with 10% normal donkey serum (Chemicon International Inc, Temecula, CA), followed by overnight incubation at 4°C with the primary antibody.

The following primary antibodies were used for immunohistochemistry: EGFP (1:500, Aves, #GFP-1020); H3K27me3 (1:200, Millipore-Sigma, #07-449); HCN4 (1:40, Alomone, #APC052); sm22a (1:500, Abcam, #ab14106); anti-sarcomeric α-acinin (1:100, Abcam, #ab9465); Tyrosine Hydroxylase (1:500, Novus Biologicals, #NB300); Isl1 (1:1000, ab109517, Abcam; or a mixture of 1:10, #40.2D6 and 1:100, #39.4D5, Developmental Studies Hybridoma Bank); cardiac myosin light chain-2 (1:200, Novus, NBP1-40754); Nkx2.5 (1:50, Santa Cruz Biotechnologies, #SC8697), cardiac troponin-T (1:200, Abcam, ab8295); Neurofilament M (1:200, Abcam, #7794). Subsequently, the antibodies were visualized by incubating the sections for 1h at 37°C with FITC, Cy3 and Cy5-conjugated F(ab’)2 fragments of affinity-purified secondary antibodies (1:200, Jackson Immunoresearch) or Alexa-fluor 488, 568 and 633 dyes (1:500, ThermoFisher).

#### Western Blot

Protein lysates were prepared in RIPA buffer and quantified with the Bradford Assay (Bio-Rad). Electrophoresis was performed in precast, NuPage 4-12% Bis-Tris protein gels beofre transferring into PVDF membranes using the Trans-Blot Turbo transfer system (Biorad). Prior to blocking and antibody incubation, membranes were stained with Ponceau S reagent (Sigma-Aldrich, #P3504) to visualize protein bands and cut at specific sizes so different antibodies could be tested at the same time. Western blots were performed according to previously described protocols (Hatzistergos et al., 2019) using antibodies against H3K27me3 (Sigma-Aldrich, rabbit polyclonal, 0.4μg/ml), Isl1 (Rabbit monoclonal, Abcam, 1:10,000), Hsp90 (Rabbit polyclonal, Cell Signaling, 1:2,000), Gapdh (Rabbit Monoclonal, Cell Signaling, 1:2000) and a Goat anti-rabbit IgG, HRP-linked antibody (Cell Signaling, 1:2,000). Densitometry analysis of western blots was performed using Fiji-Image J.

#### X-gal staining and quantification

X-gal staining was performed as previously described (Hatzistergos et al., 2015). Briefly, whole hearts were collected, washed in ice-cold Hank’s Balanced Salt solution and fixed in 4% paraformaldehyde at 4°C, for 30 minutes. Following fixation, samples were rinsed twice in ice-cold PBS and permeabilized overnight at 4°C, by shaking gently in wash solution (PBS, 0.02% sodium deoxycholate, 0.01% IGEPAL). The next day, samples were transferred in staining solution [wash solution supplemented with 5mM K3[Fe(CN)6], 5 mM K4Fe(CN)6·3H2O, 2mM MgCl_2_, and 1mg/ml x-gal] and incubated overnight at 37°C, with gentle shaking. Samples were then washed in wash solution to remove any precipitation, fixed for 10 minutes in 2% paraformaldehyde and imaged in a Zeiss, Discovery V8 stereomicroscope equipped with a Nikon D7200 digital camera. For x-gal quantification, whole-heart images in jpeg format and their respective x-gal^+^ areas were selected in Adobe Photoshop Elements (version 12.1) using the semi-automated color-range selection tool, and converted into pixel values. The amount of x-gal was expressed as the percentage of x-gal^+^ pixels per total pixels of the whole heart.

#### Quantification of myocardial contribution in Isl1*^fl/fl^*, *Wnt1/GAL4/Cre-11* and *Wnt1-Cre2* mice

Mouse hearts were fixed in 4% paraformaldehyde for 24h at 4°C, followed by overnight incubation in 30% sucrose, and embedded in OCT (TissueTek), as previously described(Hatzistergos et al., 2015). Long-axis 10μm-thick cryosections were prepared and immunostained for cardiac myosin light chain-2 (1:200, Novus, NBP1-40754) followed by a Cy-5 conjugated donkey anti-rabbit secondary antibody, as previously described (Hatzistergos et al., 2015). Slides were mounted with fluoroshield mounting medium with DAPI (Abcam, #ab104139), and imaged on a Zeiss LSM-710 confocal microscope (Carl Zeiss MicroImaging, Inc. Thornwood, NY), using the Zeiss ZEN software (version 2009, Carl Zeiss Imaging Solutions, GmbH). For *Confetti*, samples were imaged in a Leica SP5 inverted microscope at the University of Miami Analytical Imaging Core Facility. To quantify the % of ventricular myocardium expressing *Cre* reporters, the areas of Cre^+^/cmlc2^+^ were quantified and expressed as a % of the total area of Cy-5^+^ (cmlc2^+^) myocardium per slide, using the ImageJ area analysis software as previously described(Fernando et al., 2011). A total of 2-5 slides were analyzed per heart.

#### Quantitative Polymerase Chain Reaction

Gene-expression analysis was performed as described before (Hatzistergos et al., 2016; Hatzistergos et al., 2015). Briefly, following total RNA extraction with the RNeasy mini kit (Qiagen) and cDNA synthesis with the high-capacity reverse-transcription kit (Applied Biosystems), samples were subjected to quantitative polymerase chain reaction in a iQ5 real time PCR detection system (BioRad), using the Taqman Universal Master mix (Applied Biosystems). The following probes were used: *Gapdh* (Mm99999915), *Isl1* (Mm00517585_m1), *Wnt1* (Mm01300555_g1), *Nkx2-5* (Mm01309813_s1).

#### H3K27me3 and H3K27ac ChIP-Seq Analysis

The following publicly available datasets were obtained from the mouse ENCODE project (Consortium, 2012; Sloan et al., 2016). For H3K27me3 ChIP-seq mouse hearts: E10.5 (ENCSR266JQW), E14.5 (ENCFF129FZH), PN0 (ENCSR782DGO) and postnatal week 8 (ENCFF344SFV); for H3K27me3 ChIP-seq mouse hindbrain: E10.5 (ENCSR582S); For H3K27ac ChIP-seq mouse hearts: E10.5 (ENCS582SPN), E14.5 (ENCFF034YQZ), PN0 (ENCSR675HDX) and postnatal week 8 (ENCFF691YDA); for H3K27ac ChIP-seq mouse hindbrain: E10.5 (ENCSR594JGI); The mouse *Isl1* enhancer and promoter regions were obtained from the UCSC Genome Browser database (Kent et al., 2002) [UCSC Genome Browser on Mouse Dec. 2011 (GRCm38/mm10) Assembly, http://genome.ucsc.edu]. For the analysis, fold-over-control signal tracks were used, depicting control-normalized tag density from 2 biological replicates per time point, pooled together.

#### Generation, characterization and differentiation of iPSC*^Wnt1-GOF^*

The iPSC*^Wnt1-GOF^* were generated from tdTomato-negative neonatal cardiac fibroblasts, derived from *Wnt1/GAL4/Cre-11;tdTomato* mice, using a polycistronic lentivirus (STEMCCA, Millipore) as previously described (Hatzistergos et al., 2018; Hatzistergos et al., 2015). iPSC*^Wnt1-GOF^* were propagated without feeders on 0.1% gelatin-coated plates (Millipore), with NDiff227 (Clonetech), supplemented with 1000 units/ml LIF (Millipore), 1µMPD0325901 (Tocris) and 3µM CHIR99021 (Tocris). For CNC differentiation, iPSC*^Wnt1-GOF^* were trypsinized into single cells (Thermo), washed twice, and resuspended at a final concentration of 25,000 cells/ml in IMDM, 2mM L-glutamine, 20% FBS (Thermo), 0.1mM non-essential aminoacids and 0.1mM β-mercaptoethanol (all from Thermo). Embryoid body differentiation was performed using the hanging-drop method, as previously described (Hatzistergos et al., 2018; Hatzistergos et al., 2015). During the first 48h of differentiation, EBs were supplemented with 2μM Dorsomorphin (Tocris) or DMSO. For canonical Wnt/β-catenin inactivation, EBs were supplemented with 1μM XAV939 or DMSO on day 9 of differentiation. For canonical Wnt/β-catenin activation EBs were supplemented with 3μΜ CHIR99021 or DMSO on day 9 of differentiation. All samples were cultured under 5% CO2 atmosphere at 37°C. EBs were monitored daily for the emergence and quantification of tdTomato^+^ beating cells, in an Olympus IX81 fluorescent microscope as described before(Hatzistergos et al., 2018; Hatzistergos et al., 2016; Hatzistergos et al., 2015).

### Isolation of primary cardiac cells for single-cell RNA-seq and in-vitro pharmacologic experiments

For the scRNAseq and MG132 inhibitor experiments 3-day old *Isl1^+/+^* male and female neonatal mice were used (n=15 for each experiment). Briefly, the cardiac apex as well as the dorsal and ventral domains of the cardiac base, were dissected from each neonate and combined together to generate 3 pools of apical, dorsal and ventral cardiac tissues, respectively. The ventral domain included the proximal aorta, pulmonary artery, and OFT base. The dorsal domain included the SAN and CGs. Each pool was subsequently washed twice in D-PBS supplemented with 1% penicillin-streptomycin, and minced into ∼2mm fragments as previously described (Hatzistergos et al., 2018; Hatzistergos et al., 2016; Hatzistergos et al., 2015). Samples were then washed again, re-suspended in 3ml digestion buffer (Hanks balanced salt solution, supplemented with 2mg/ml collagenase type IV and 1.2U/ml Dispase II and 1% penicillin/streptomycin) and incubated at 37°C for 15 min with gentle rocking. Digested samples were then gently triturated 10-20 times using a 10 ml serological pipette. This process was repeated 3 more times, for a total digestion time of 45 min. Dissociated cells were then collected and washed 2 times with D-PBS, by centrifuging at 100xg for 3min. Finally, the cell suspension was passed through a 40μm nylon-strainer, resuspended in plating medium ([RPMI1640, supplemented with 1% L-glutamine, 1% penicillin/streptomycin and 2% B27 minus insulin) and seeded in 0.1% gelatin-coated, 12-well or 24-well plates, at a density of 100,000 cells/ml. Cells were fed with fresh plating medium every other day for a total of 5 days. For the MG132 experiments, 4 wells of a 24-well plate from each cell pool were cultured on day 5 with or without 25μM MG132, for 90 min. For the scRNAseq experiment, 4 wells of a 24-well plate from the dorsal and ventral pools were dissociated on day 5 with Tryple (ThermoFisher Scientific), passed through a 40μm nylon-strainer to ensure single-cell suspensions, and processed immediately for library preparation. For the EED226 experiments, 4-week old *Isl1-nLacZ* male and female mice were used (n=4), and each sample was cultured separately (in triplicates) for 5 days, in the presence or absence of 10μM EED226.

### Single-Cell RNA-seq Analysis

The dorsal and ventral single-cell RNA-Seq libraries were prepared with the 10X Chromium system, using the Chromium Single Cell 3’ (v3 chemistry) kit, according to the manufacturers’ instructions, and sequenced in an Illumina Novaseq 6000 at the University of Miami, Sylvester Comprehensive Cancer Center, Oncogenomics Core Facility. Raw base call (BCL) files were analyzed using CellRanger. The “mkfastq” command was used to generate FASTQ files and the “count” command was used to generate raw gene-barcode matrices aligned to the Mm10 Ensembl build 93 genome. The data from both samples were combined in R using the Seurat package and an aggregate Seurat object was generated (Butler et al., 2018; Stuart et al., 2018). To ensure our analysis was on high-quality cells, filtering was conducted by retaining cells that had unique molecular identifiers (UMIs) greater than 400, expressed 100 to 8000 genes. This resulted in 9,893 cells for Dorsal and 11,418 cells for Ventral. Data from each sample were analyzed using the SCTransform Integration Workflow (https://satijalab.org/seurat/v3.0/integration.html). Data from each sample were normalized using the SCTransform() function and integration features were identified using SelectIntegrationFeatures() with “nfeatures” and “fvf.nfeatures” set to 5000 (Hafemeister and Satija, 2019). To identify integration anchor genes among the 2 samples the PrepSCTIntegration() and FindIntegrationAnchors() functions were used with 5000 genes. Using Seurat’s IntegrateData() the samples were combined into one object. To reduce dimensionality of this dataset, principle component analysis (PCA) was used and the first 10 principle components further summarized using uniform manifold approximation and projection (UMAP) dimensionality reduction (McInnes et al., 2018). The DimPlot() function was used to generate the UMAP plots displayed. Clustering was conducted using the FindNeighbors() and FindClusters() functions using the original Louvain algorithm (Blondel et al., 2008), 10 PCA components, and a resolution parameter set to 0.5. The resulting 15 Louvain clusters were visualized in a two-dimensional UMAP representation and were annotated to known biological cell types using canonical marker genes, as well as gene set enrichment analysis (Enrichr) (Kuleshov et al., 2016). The DoHeatmap() command was used to generate the heatmap displayed and the VlnPlot() command was used to generate violin plots of the Louvain clusters. Functional enrichment analysis of the differentially expressed genes between the ventral and dorsal *Isl1^+^* subsets was conducted with G:Profiler (Raudvere et al., 2019) and GSEA (Mootha et al., 2003).

### Single-Cell Trajectory Reconstruction

Single-cell pseudotime trajectories were constructed with Monocle 2 (Qiu et al., 2017b; Trapnell et al., 2014). For the analysis we utilized the normalized expression data from the subset expressing *Isl1* or *Nkx2-5* greater than 0 to infer the relationship between these cell types. Using these criteria, 868 cells entered the Monocle2 analysis. Genes for trajectory inference were selected using the dispersionTable() function to calculate a smooth function describing how variance in each gene’s expression across cells varies according to the mean. 5,051 genes with mean expression greater than or equal to 0.1 were used for the analysis. The reduceDimension() function was utilized with the DDRTree (Qiu et al., 2017b) reduction method and the following parameters modified: ncenter = 100, max_components = 3, and num_dim = 10. Results were visualized using the plot_cell_trajectory() or plot_complex_cell_trajectory function and annotated with the corresponding cell type. BEAM was used for branched-dependent gene expression analysis (Qiu et al., 2017a). Functional enrichment analysis was conducted with G:Profiler (Raudvere et al., 2019) and GSEA (Mootha et al., 2003).

### Statistical analysis

The scRNAseq data were analyzed in Rstudio using the Seurat (Stuart et al., 2019) and Monocle (Qiu et al., 2017a) libraries, as described above. For differential expression analysis in Seurat, the default two-sided non-parametric Wilcoxon rank sum test was utilized. Trajectory-dependent differential expression analysis, was conducted with Monocle’s Branched expression analysis modeling (BEAM). Functional enrichment analysis of the differentially expressed genes between the ventral and dorsal *Isl1^+^* subsets was conducted with G:Profiler (Raudvere et al., 2019) and GSEA (Mootha et al., 2003). All other statistical tests were performed in GraphPad Prism (Version 8, La Jolla, CA) using Student’s t-test, Mann-Whitney, or one-way ANOVA followed by Tukey’s post-hoc tests. All data met the assumptions of the tests (Bartlett’s test for normality). A p<0.05 was considered statistically significant. All values are reported as mean±SEM.

## DATA AND CODE AVAILABILITY

Next-generation sequencing data will be made available to the Gene Expression Omnibus. All custom code is available from the authors upon request.

## Supplemental Information

### Supplemental tables

**Table S1. scRNAseq cluster-specific genes. Related to Fig. 1.**

**Table S2. Differentially expressed genes between the dorsal and ventral *Isl1^+^* scRNAseq subsets. Related to Fig. 2.**

**Table S3. g:Profiler functional enrichment analysis of the differentially expressed genes from **Table S2** (q-val< 0.05). Related to Figs. 2-3.**

**Table S4. GSEA functional enrichment analysis of the differentially expressed genes from **Table S2** (q-val< 0.05). Related to Fig. 2.**

**Table S5. Differentially expressed genes based on BEAM analysis on branch-point 1. Related to Fig. 5.**

**Table S6. g:Profiler functional enrichment analysis of the differentially expressed genes from **Table S5** (q-val< 0.01). Related to Fig. 5.**

### Supplemental Figure Legends

**Figure S1.**
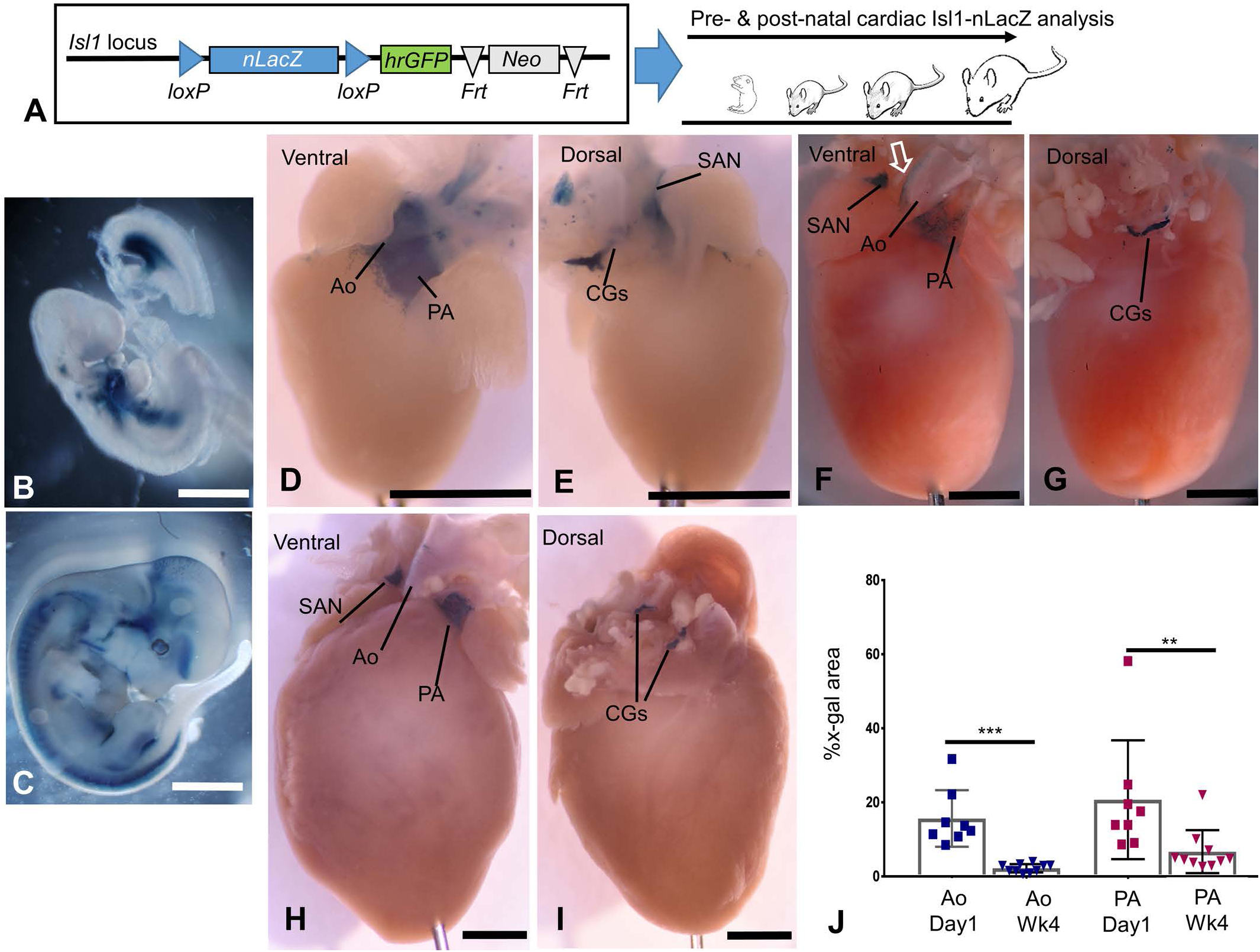
Temporal and spatial analysis of cardiac *Isl1* expression under the *Isl1-nLacZ* knock-in. **A**. Illustration of the allele and experimental outline. **B-C**, X-gal stained embryos at E9.5 (B) and E12.5 (C). **D-I**, Ventral and dorsal views of cardiac x-gal stained heart at PN1 (D-E), PN week 2 (F-G) and PN100 (H-I). **J**, Quantification of the x-gal+ areas as a percentage of the total area of the heart at PN1 (n=8 mice) and week (wk) 4 (n=10 mice). PA, pulmonary artery; Ao, aorta; CGs, cardiac ganglia; SAN, sinoatrial node. ***p<0.0001, and **p=0.0204 (unpaired t-tests). Values are mean±SEM. Scale bars, 1mm. Related to Figs. 1-7.

**Figure S2.**
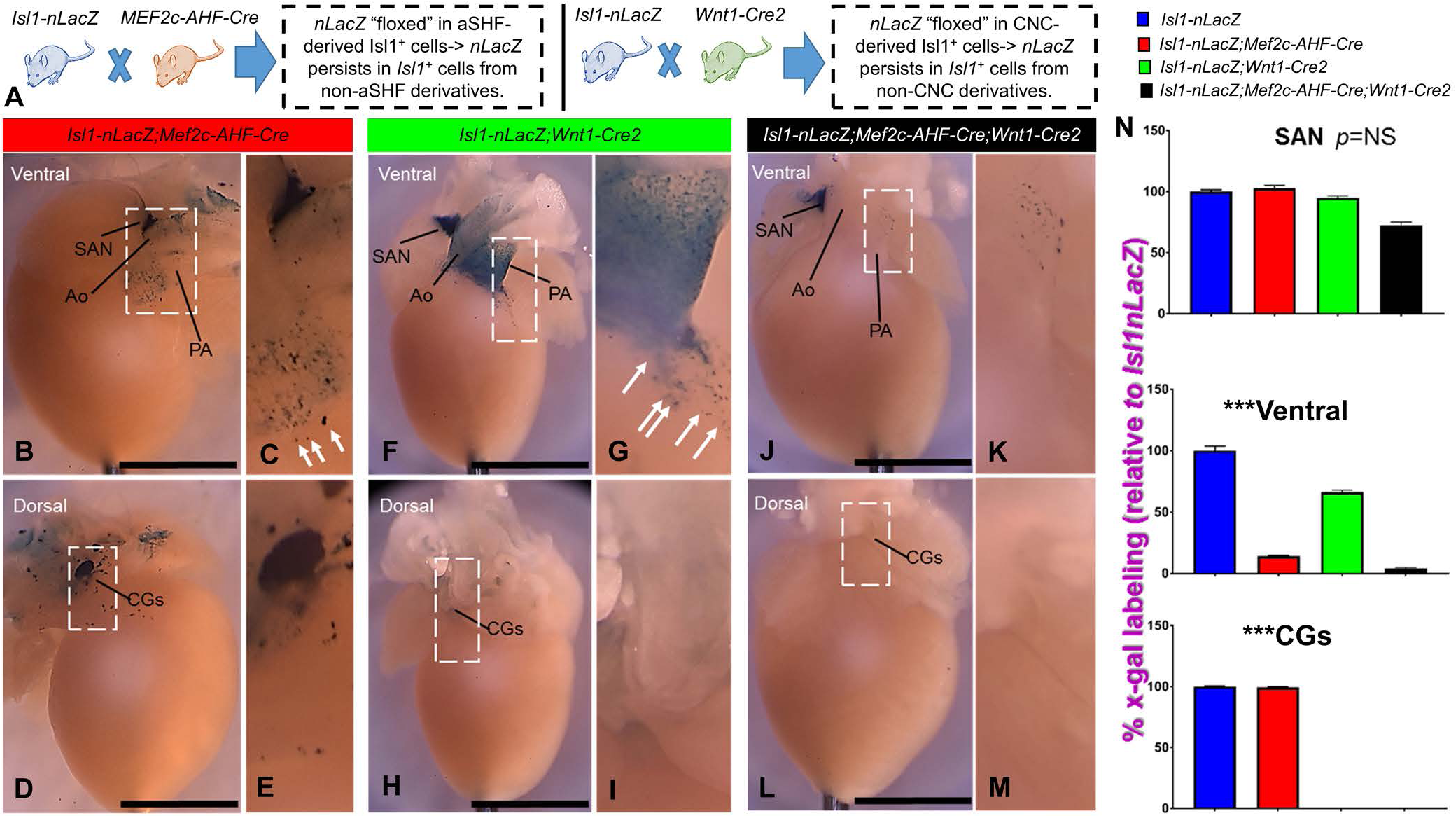
Lineage-tracing of postnatal *Isl1-nLacZ* activity. **A**, Experimental outline of ventral and dorsal *Isl1-nLacZ* lineage-tracing. The *Isl1-nLacZ* reporter is conditionally excised from anterior heart field (aSHF) and neural crest (CNC) derivatives under the *Wnt1-Cre2* and *Mef2c-AHF-Cre* mouse lines, respectively. **B-E**, At postnatal day 1 (PN1), *Isl1-nLacZ;Mef2c-AHF-Cre* hearts display strong dorsal x-gal staining in the sinoatrial node (SAN, B) and cardiac ganglia (CGs, D, inset; E, higher magnification). Ventrally, staining in the pulmonary artery (PA) and aorta (Ao) is reduced, but not extinguished (B, inset; C, higher magnification). Importantly, a few x-gal+ myocardial cells are present in the outflow tract (OFT) base (C, arrows). **F-I**, In *Isl1-nLacZ;Wnt1-Cre2* hearts, dorsal x-gal staining is sustained in the SAN (F) but extinguished from CGs (H, inset; I, higher magnification). Ventrally, x-gal is sustained in the Ao, PA (F) and OFT base (F, inset; G, higher magnification, arrows). **J-M**, In triple-mutant *Isl1-nLacZ;Mef2c-AHF-Cre;Wnt1-Cre2* hearts, dorsal x-gal staining is sustained in the SAN (J) but extinguished from CGs (L, inset; M higher magnification). Ventrally, a few x-gal+ cells remain scattered in the proximal PA and dorsolateral Ao wall (J, inset; K, higher magnification), indicating they originate from outside the *Wnt1-Cre2* and *Mef2c-AHF-Cre* lineages. Scale bars, 1mm. n= 5 *Isl1-nLacZ*; n= 10 *Isl1-nLacZ;Mef2c-AHF-Cre; n= 6 Isl1-nLacZ;Wnt1-Cre2; and n= 4 Isl1-nLacZ;Mef2c-AHF-Cre;Wnt1-Cre2* mice. ***p<0.0005 (Kruskal-Wallis with Dunn’s multiple comparison test). Values are mean±SEM. Scale bars, 1mm. Related to Figs. 1-7.

**Figure S3.**
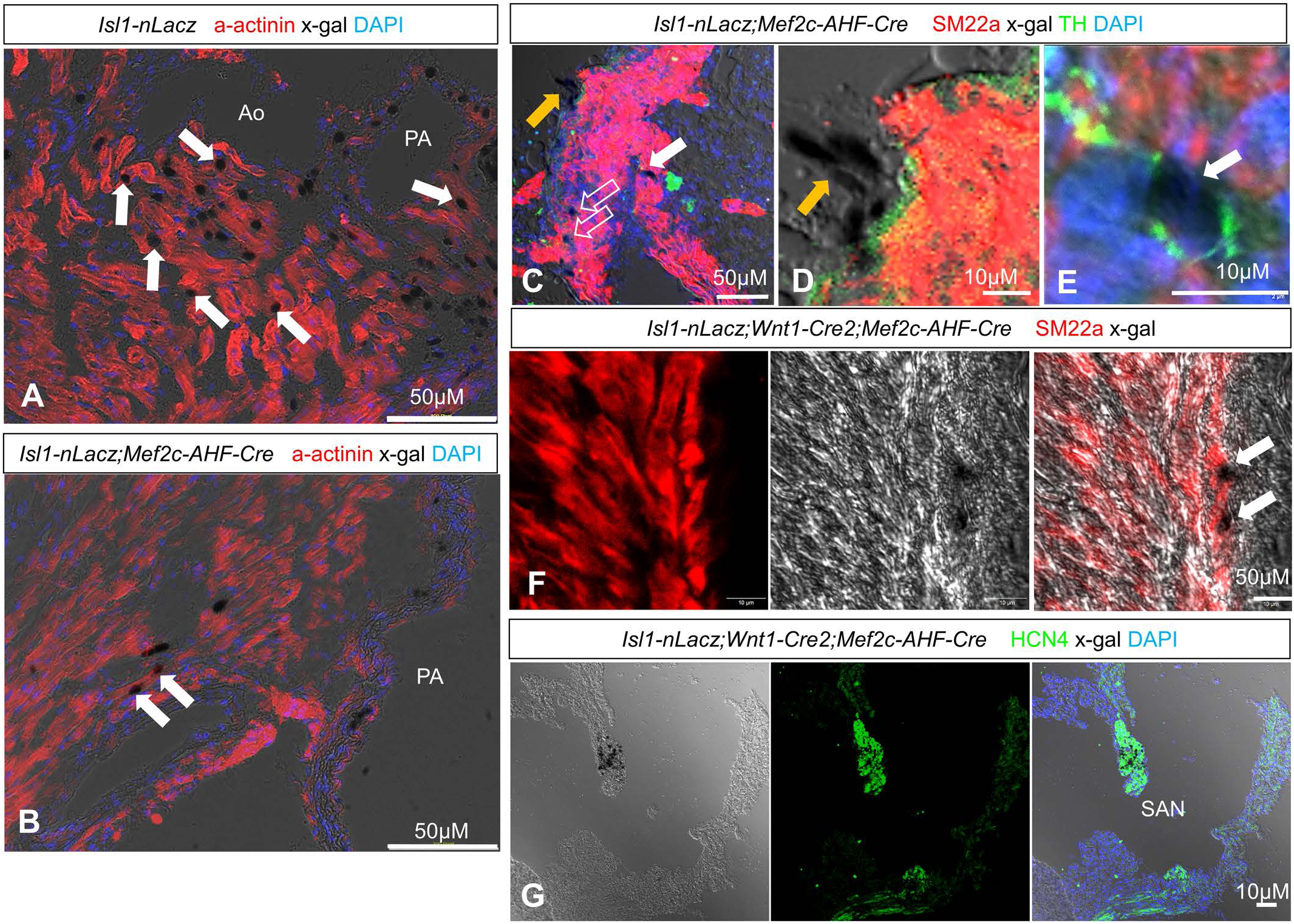
Characterization of postnatal *Isl1-nLacZ*^+^ cardiac cells. **A-B**, co-localization of x-gal with α-sarcomeric actinin^+^ cardiomyocytes (arrows) within the OFT base myocardium of *Isl1-nLacZ* (A) and *Isl1-nLacZ;Mef2c-AHF-Cre* (B) neonatal mice. **C-E**, co-localization of x-gal with sm22a^+^ smooth muscle cells (C, open arrows) and tyrosine hydroxylase (TH)^+^ sympathetic nerves (C, E white arrows) in *Isl1-nLacZ;Mef2c-AHF-Cre* neonatal mice. X-gal+ cells which do not co-localize with either marker are also shown (C-D yellow arrows). **F-G**, co-localization of x-gal with sm22a^+^ smooth muscle cells (F, arrows) and HCN^+^ sinoatrial node (SAN) cells (G, arrows) in *Isl1-nLacz;Wnt1-Cre2;Mef2c-AHF-Cre* neonatal mice. n= 5 Isl1-nLacZ; n= 10 *Isl1-nLacZ;Mef2c-AHF-Cre; n= 6 Isl1-nLacZ;Wnt1-Cre2; and n= 4 Isl1-nLacZ;Mef2c-AHF-Cre;Wnt1-Cre2* mice. Scale bars are depicted within each figure panel. Related to Figs. 1-7.

**Figure S4.**
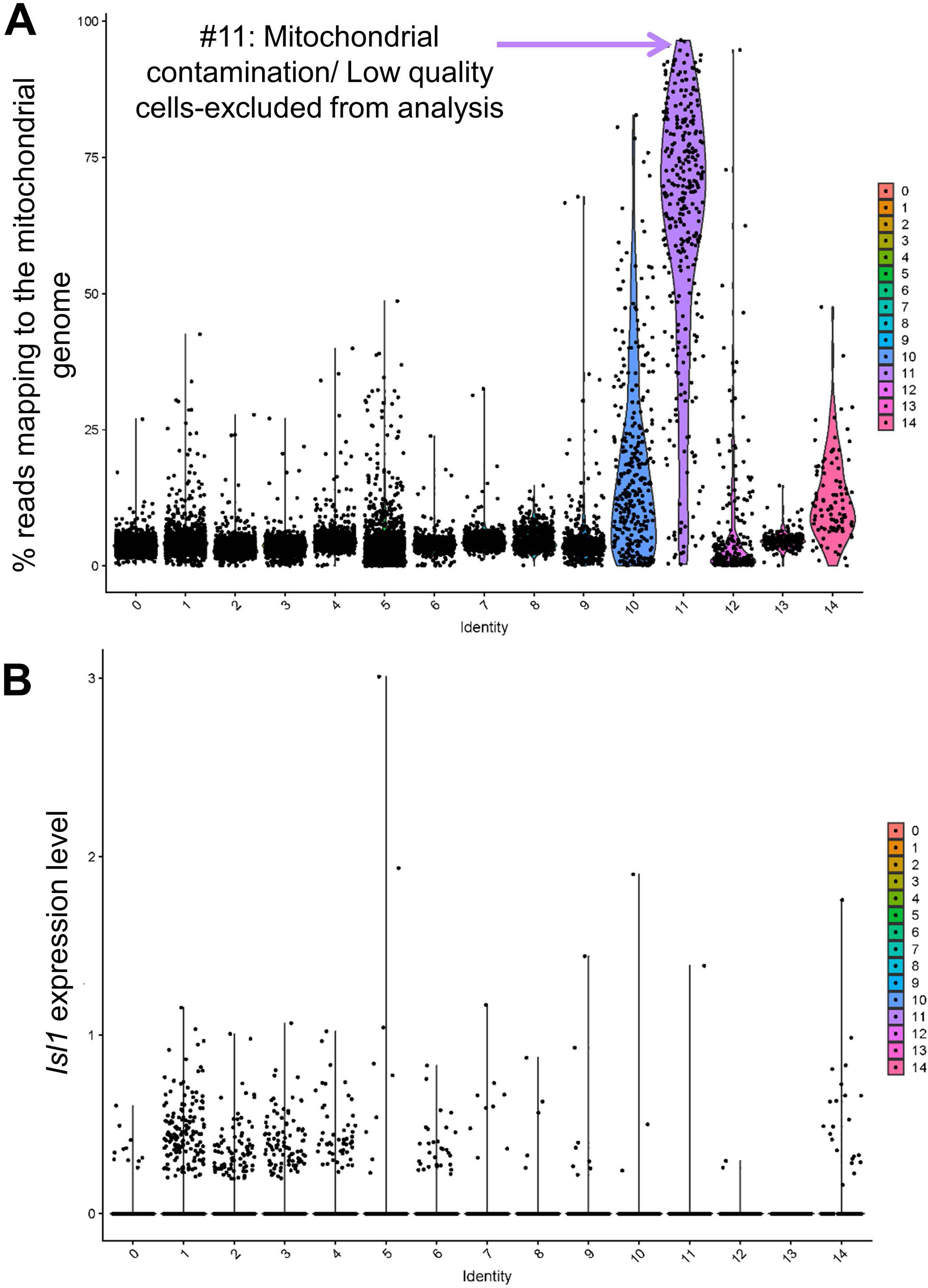
scRNAseq of arterial (ventral) and venous (dorsal) pole postnatal domains. **A,** Violin plot of the percentage of mitochondrial (mt) genes present within each UMAP cluster. **B**, Violin plot visualization of Isl1 expression within each cluster. Related to Fig. 1.

**Figure S5.**
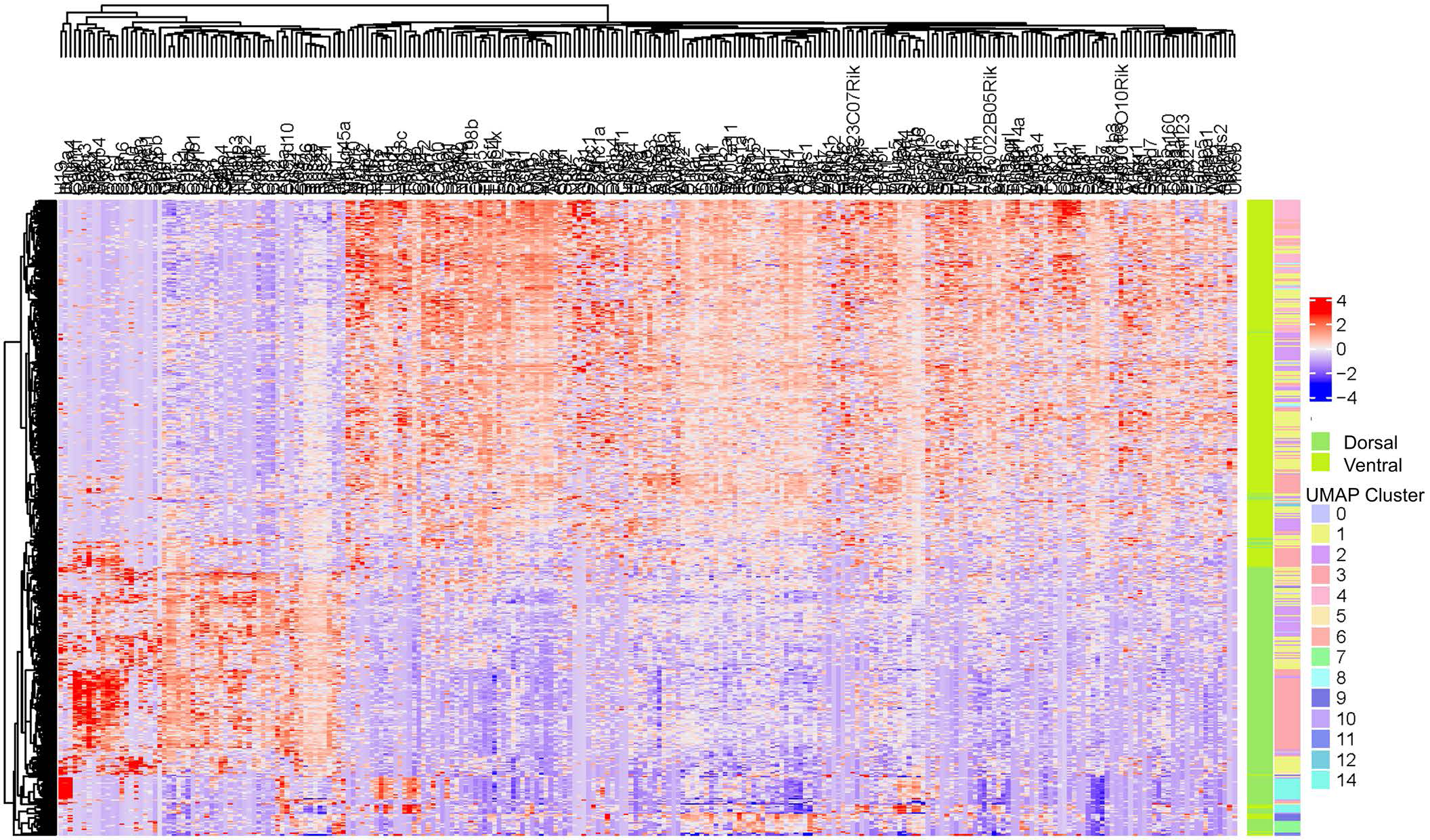
Differentially expressed genes (DEGs) of dorsal vs ventral *Isl1^+^* scRNAseq subsets. Unsupervised hierarchical clustering of the top 250 DEGs based on their dorsoventral and UMAP cluster identities. Related to Fig. 2.

**Figure S6.**
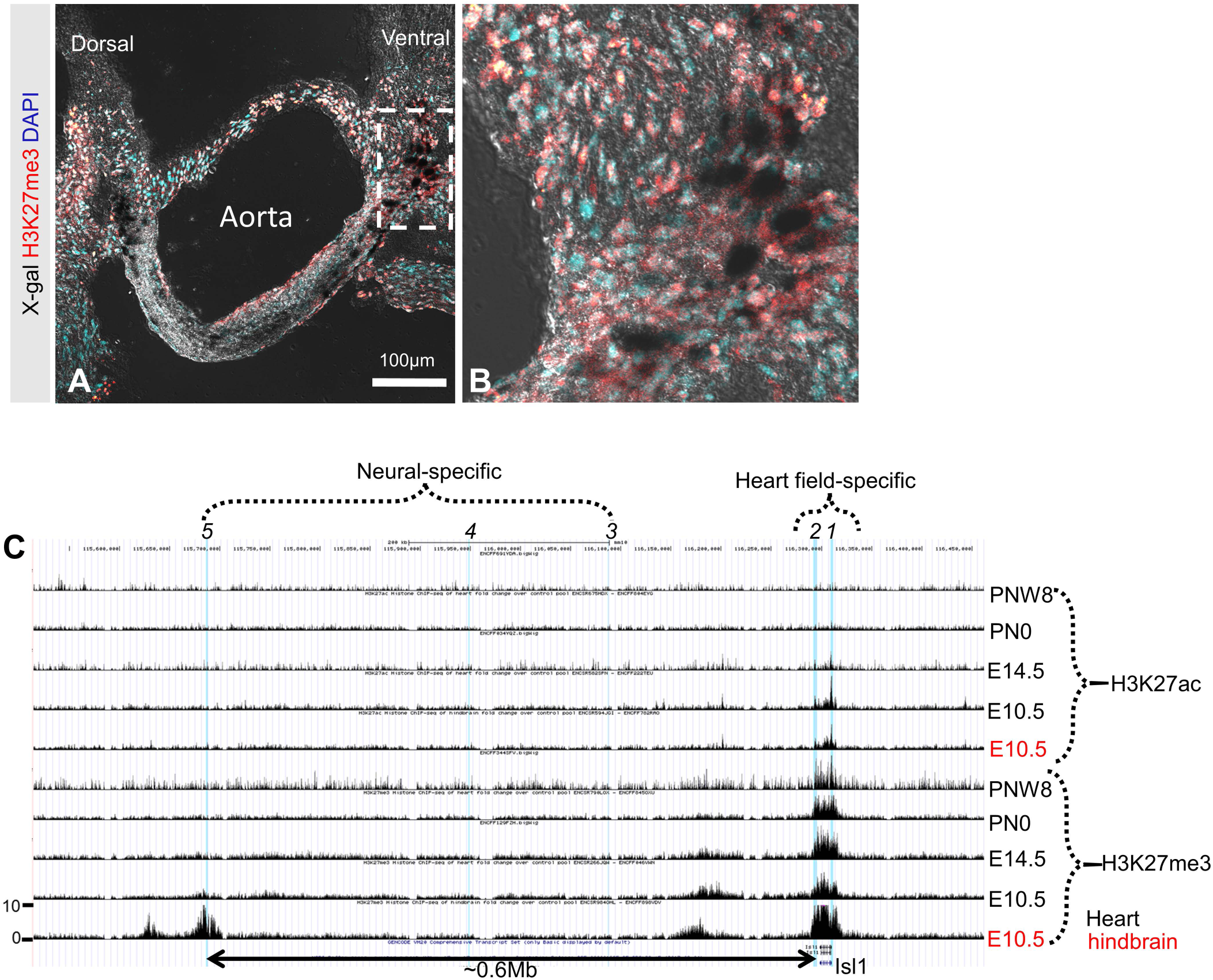
H3K27me3-mediated silencing of heart field (ventral) *Isl1* expression. **A-B**, x-gal^+^ cells in the aorta of a PN-week *4 Isl1-nLacZ* heart co-localize with H3K27me3. Panel B is a higher magnification of the boxed region in panel A. **C**, H3K27me3 and H3K27ac ChIP-seq tracks from embryonic day (E) 10.5, 14.5, postnatal day (PN) 0 and postnatal week-8 mouse hearts (black). For comparison, E10.5 hindbrain tracks are also shown in red. The following RefSeq functional elements are highlighted in blue: *1*, Isl1 promoter fragments; *2*, Isl1-F and Isl1-F2 heart field-specific enhancers; *3*, E1 enhancers in motor neurons and hindbrain; *4*, E2 enhancer in motor neurons, induced by Onecut1; and *5*, VISTA enhancer mm933 with predicted enhancer activity in dorsal root ganglion, cranial nerve and nose. Note that the transcriptional repressor H3K27me3 accumulates in the promoter and heart field-specific elements as early as E10.5. In addition, accumulation of the transcriptional activator H3K27ac is virtually diminished by E14.5. In contrast, all neural-specific Isl1 regulatory elements are H3K27me3-free. Values represent fold-over-control signal tracks from 2 biological replicates per time point. Scale bar, 100µm. Histone tracks are visualized on UCSC Genome Browser on Mouse Dec. 2011 (GRCm38/mm10) Assembly.

**Figure S7.**
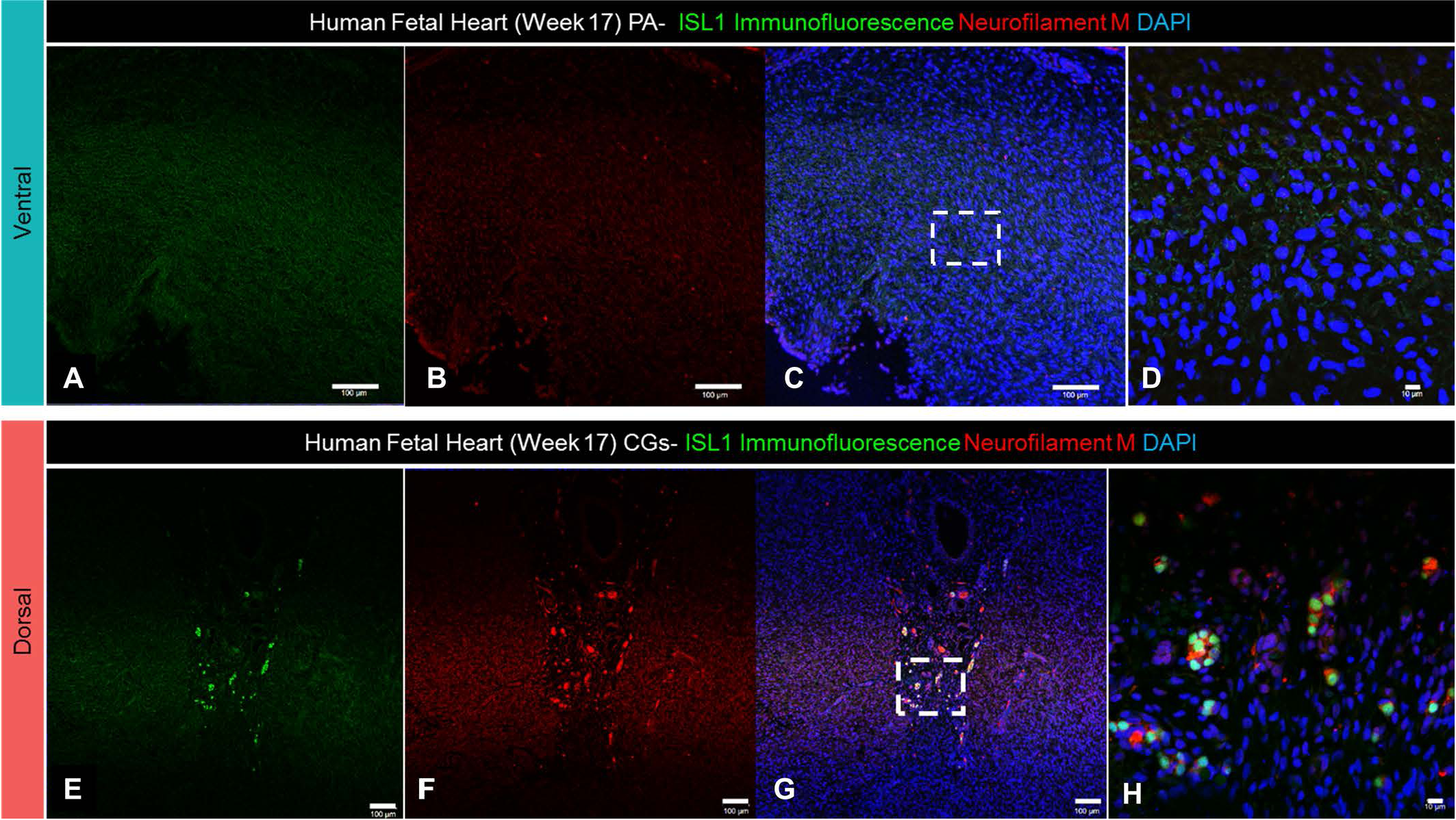
Degradation of ventral, but not dorsal, *ISL1* expression in the developing human heart. **A-D.** Lack of ISL1 immunoreactivity in the proximal pulmonary artery of the developing human heart. **E-H**, Co-localization of ISL1 with neurofilament M in the venous pole domain indicates that ISL1 immunoreactivity identifies cardiac neural crest-derived ganglia. Panels D-H are high magnifications of the boxed areas in C and G, respectively. Scale bars, 100µm (A-C, E-G); 10µm (D, H). (n=2). Related to Fig. 3.

**Figure S8.**
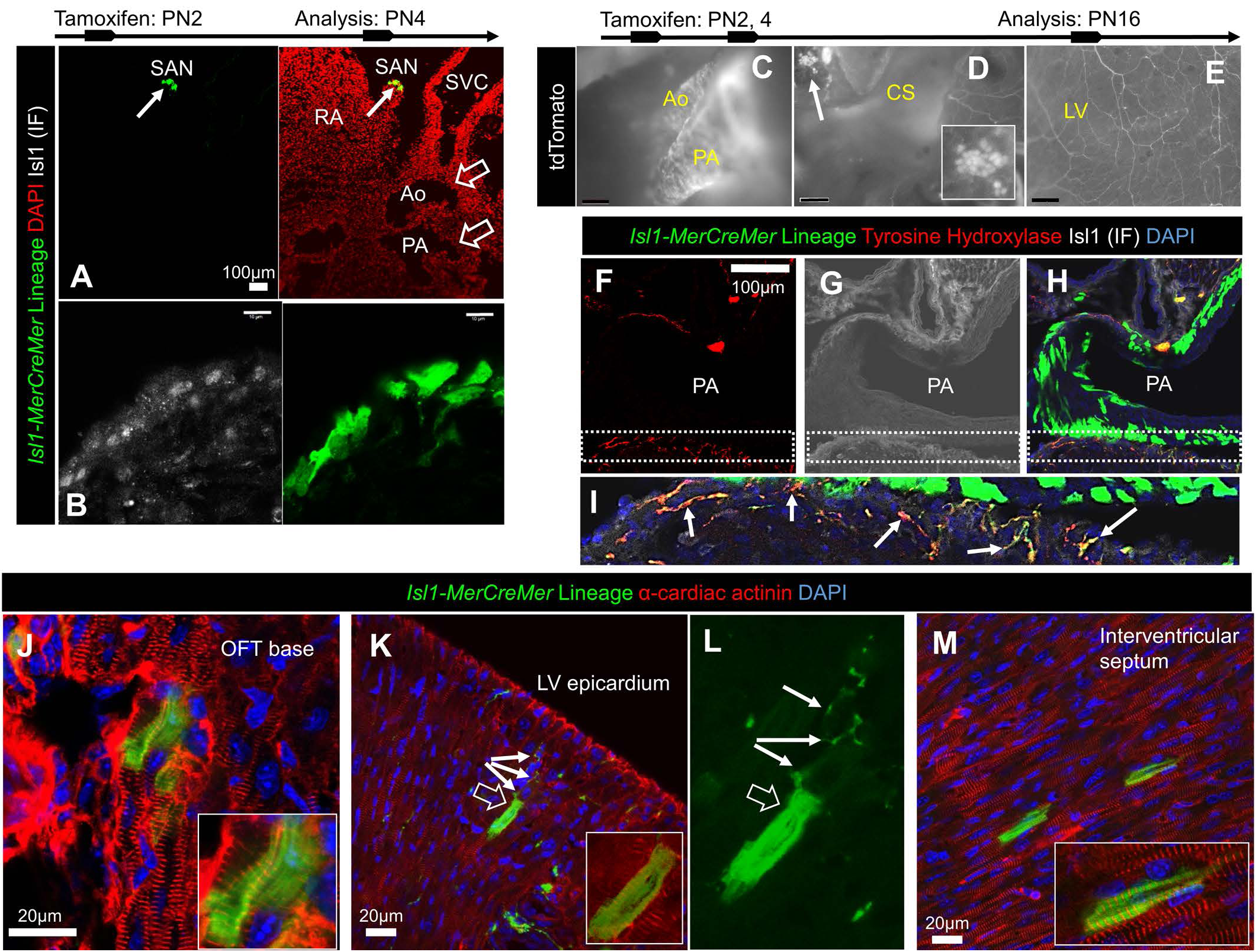
*Isl1-MCM* fate-mapping indicates extensive neurogenic and limited cardiomyogenic contribution of postnatal *Isl1*^+^ cardiac cells. **A-B**, Expression of Isl1-MCM reporters in postnatal day (PN) 4 mice (n=4) following a single dose of tamoxifen on day 2 indicates partial recombination in venous (Sinoatrial node; A, arrow; and B in higher magnification) but not arterial pole derivatives (A, open arrows). Note that, under these experimental conditions, Isl1-MCM recombination faithfully recapitulates Isl1 immunoreactivity (B). **C-D**, Live tdTomato imaging following repeated induction with tamoxifen on PN2 and PN4, and extension of fate-mapping window to 2 weeks (n=3). Tdtomato successfully marks both the arterial (C) and venous pole (D) derivatives. **E**, Live imaging of tdTomato expression in CNC-derived sympathetic nerves. **F-I**, Co-localization of Isl1-MCM;tdTomato derivatives with the sympathetic nerve marker Tyrosine Hydroxylase (I, arrows; and F, H), but not Isl1 (G, H, I), in proximal OFT derivatives. Panel I depicts a high magnification of the boxed areas in F-H. **J,** Similar to *Isl-1nLacZ*, Isl1-MCM recombination marks α-cardiac actinin^+^ myocardial cells in the proximal OFT base. **K-M**, In addition, Isl1-MCM labeling extends into rare trabecular, compact (K, open arrow and inset) and interventricular septum cardiomyocytes (M). Note the proximity of Isl1-MCM-derived cardiomyocytes and sympathetic nerves (K, L, filled arrows). Ao, aorta; PA, pulmonary artery; LV, left ventricle; OFT, outflow tract; CS, coronary sinus. Scale bars, 100μM (A, C-E, F-H); 10μM (B) and 20μM (J-M). Related to Fig. 4.

**Figure S9.**
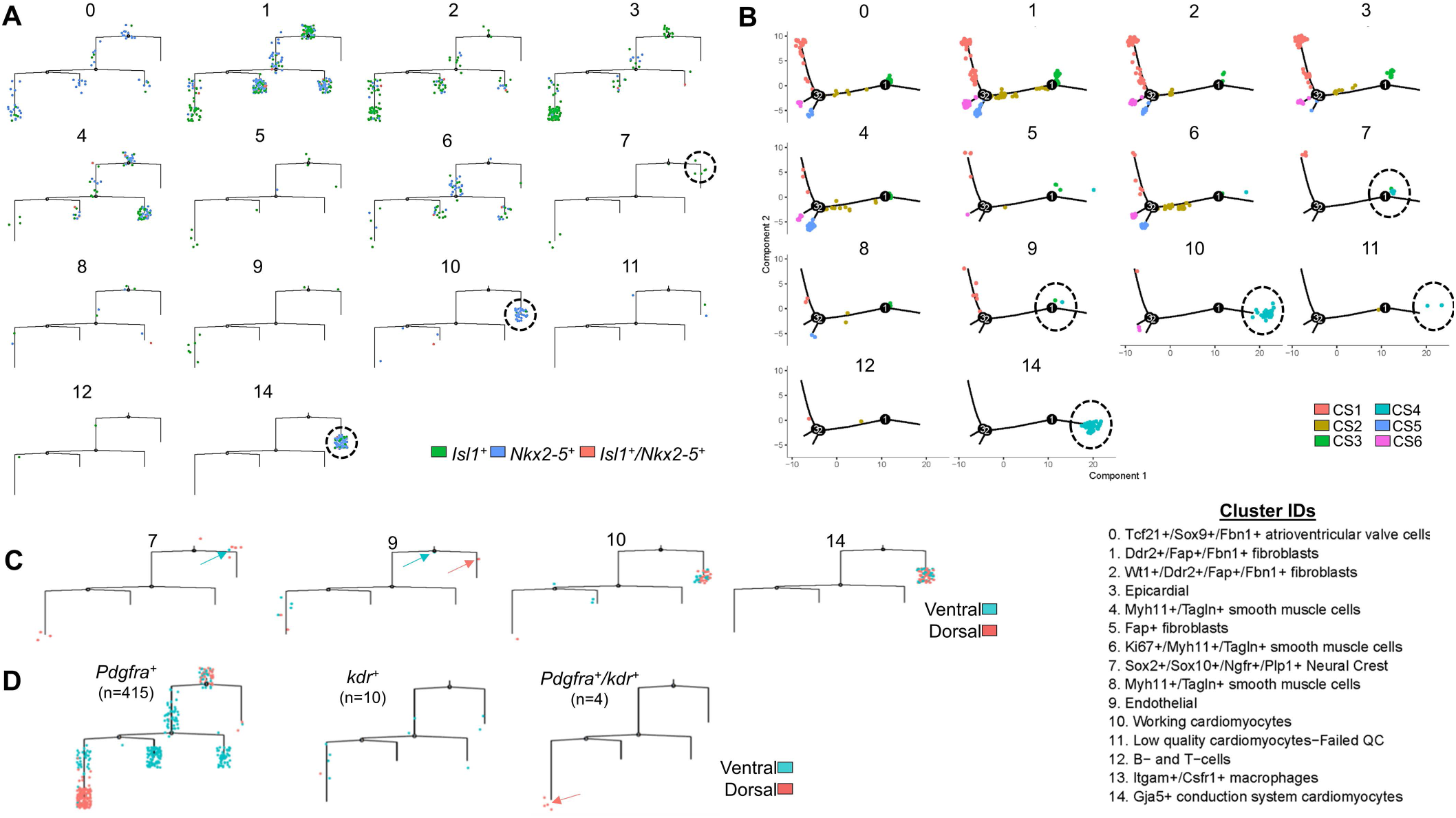
Reconstruction of pseudotime differentiation trajectory of postnatal *Isl1^+^* and *Nkx2-5^+^* lineages. **A,** Ordering of the *Isl1^+^* (green) *Nkx2-5^+^*(blue) and *Isl1^+^*/*Nkx2-5^+^* (red) subsets, based on their UMAP cluster identities. Branched trajectories are plotted as a two-dimensional tree layout. Circles delineate the *Isl1^+^* neural crest cells (cluster #7) in the myocardial trajectory; as well as the *Nkx2-5^+^* and *Isl1^+^*/*Nkx2-5^+^* cells in the myocardial clusters #10 and #14. **B,** Ordering of cells on single-cell trajectory based on their based on their UMAP cluster identities, and color-coded by cell state (CS). Circles delineate cells in clusters 7, 9, 10, 11 and 14 which belong to CS4 (myocardial state). **C**, Ordering of cells in clusters 7, 9, 10 and 14, based on their dorsoventral identities. All except one (blue arrow) *Isl1^+^* neural crest cells (cluster 7) in the myocardial trajectory are of dorsal identity, indicating functional Isl1 expression. Similarly, in the endothelial cluster (#9), there is one *Isl1^+^* ventral cell close to the root state (blue arrow) and one *Isl1^+^* dorsal cell in the myocardial trajectory (red arrow), indicating that this putative myocardiogenic cell is not aSHF-derived. **D**, Distribution of *Pdgfra^+^, Kdr^+^* and putative heart field-derived cardioblasts (*Kdr^+^/Pdgfra^+^*) in the trajectory. There are only 4 *Kdr^+^/Pdgfra^+^* cells (red arrow), all of which are in the non myocardial CS1, and are of dorsal identity, indicating that these do not represent aSHF-derived myocardial progenitors. Related to Fig. 9.

**Figure S10.**
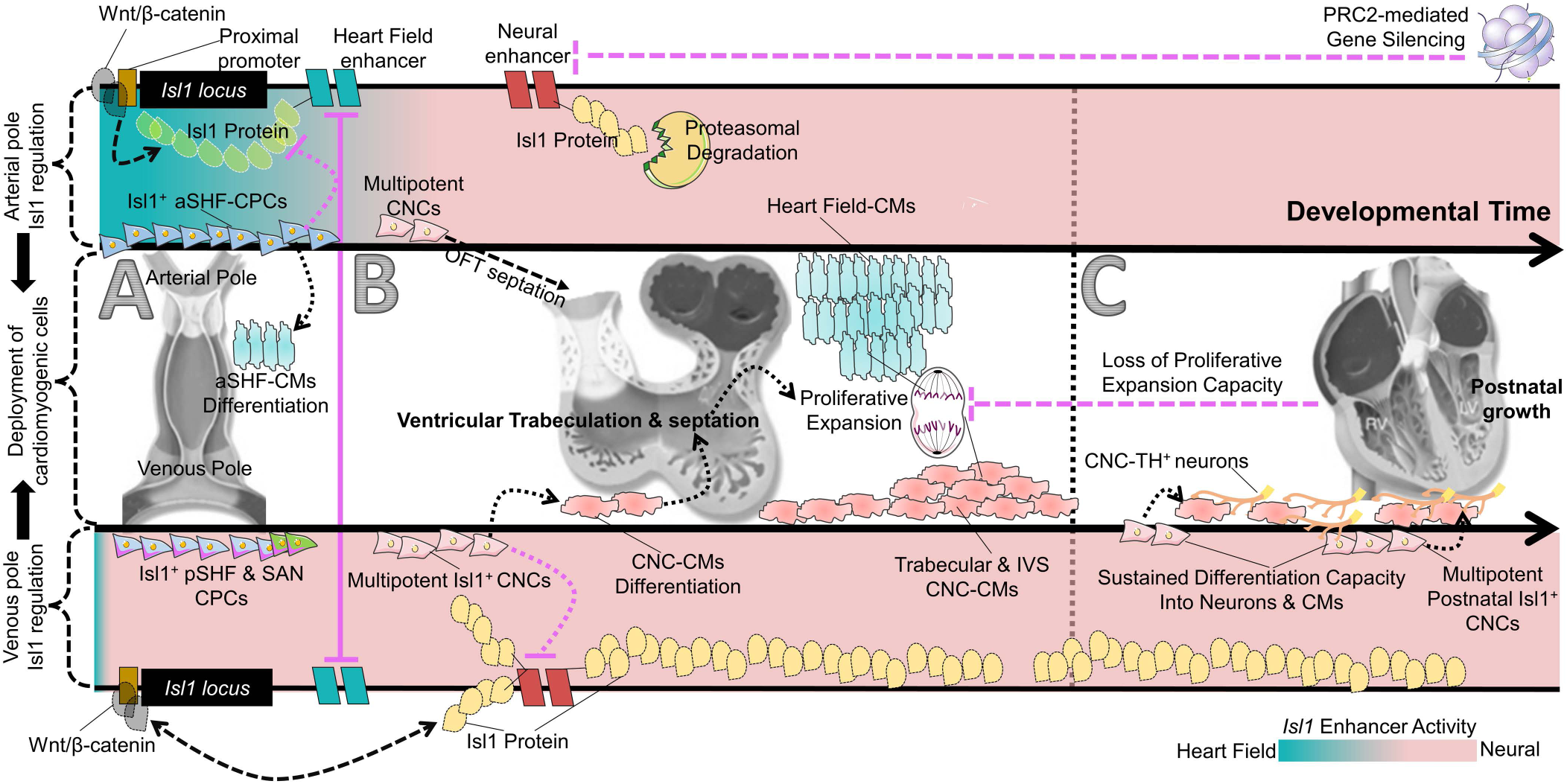
Proposed model on the role of Isl1 in embryonic and postnatal cardiac progenitors. **A.** During early cardiogenesis, Isl1 heart field (teal) and neural (rose) enhancer-specific expression promotes the recruitment of aSHF and pSHF/SAN progenitors (CPCs) through the arterial and venous poles, respectively. Repression of aSHF enhancer activity (Kappen and Salbaum, 2009), as well as activation of the miRNA17∼92 (Wang et al., 2010) and ubiquitin/proteasome pathways (Caputo et al., 2015), co-operatively promote Isl1 silencing and terminal cardiomyocyte (CM) differentiation of aSHF-CPCs. **B**, After cardiac looping, sustained neural enhancer-specific expression of *Isl1* promotes recruitment of cardiac neural crest cells (CNC) through the aSHF-derived outflow tract (OFT) and pSHF-derived dorsal mesocardium. Tissue-specific activation of the ubiquitin/proteasome pathway in the proximal OFT prevents proper Isl1 expression and, consequently, non-CM differentiation of CNCs in OFT septal derivatives. In contrast, proper dorsal Isl1 expression, sustained through a forward reinforcing loop with Wnt/β-catenin, endows CNCs with multipotent neuromuscular differentiation capacity. Dorsal CNCs migrate into the trabecular and interventricular myocardium and undergo CM differentiation in response to Wnt/β-catenin inhibition. Subsequently, CNC-CMs undergo proliferative expansion. **C**, After birth, dorsal CNCs are recruited as the principal source of postnatal cardiac sympathetic nervous system; and to a limited extent as a source of ventricular CMs, which however, do not undergo proliferative expansion after differentiation.

### SUPPLEMENTAL MOVIE LEGENDS

**Movie S1**. Live tdTomato imaging of a PN16 Isl1-MCM;TdTomato heart, following tamoxifen induction on PN2 and PN4 (n=3). Tdtomato successfully marks beating cardiomyocytes and CNC-derived sympathetic nerves, in close proximity to each other. Related to Figs. 4 and S8.

**Movie S2**. Representative movie of iPSC*^Wnt1-GOF^*-derived beating cardiomyocytes, in response to treatment with CHIR99021. The movie switches between bright-field and tdTomato epifluorescence. tdTomato+ CNC derivatives do not differentiate into cardiomyocytes. Related to Fig. 6.

**Movie S3**. Representative movie of iPSC*^Wnt1-GOF^*-derived, tdTomato^+^ beating cardiomyocytes, following transient antagonism of Wnt/β-catenin with XAV939. Panels A-C demonstrate co-expression of tdTomato with the cardiac transcription factor Nkx2.5. Related to Fig. 6.

